# The human MRS2 magnesium binding domain is a regulatory feedback switch for channel activity

**DOI:** 10.1101/2022.09.27.509701

**Authors:** Sukanthathulse Uthayabalan, Neelanjan Vishnu, Muniswamy Madesh, Peter B. Stathopulos

## Abstract

Mitochondrial RNA splicing protein 2 (MRS2) forms a major magnesium (Mg^2+^) entry channel into the matrix. While MRS2 contains two transmembrane domains that constitute a pore, most of the protein resides within the matrix. Yet, the precise structural and functional role of this obtrusive amino terminal domain (NTD) in human MRS2 function is unknown. Here, we show that the MRS2 NTD self-associates into a homodimer, contrasting the pentameric assembly of CorA, an orthologous bacterial channel. Mg^2+^ and calcium suppress lower and higher order oligomerization of MRS2 NTD, while cobalt has no effect on the NTD but disassembles full-length MRS2. Mutating pinpointed residues mediating Mg^2+^ binding to the NTD, not only selectively decreases Mg^2+^ binding affinity ∼7-fold but also abrogates Mg^2+^ binding-induced changes in secondary, tertiary and quaternary structures. Disruption of NTD Mg^2+^ binding strikingly potentiates mitochondrial Mg^2+^ uptake in wild-type and Mrs2 knockout cells. Our work exposes a mechanism for human MRS2 autoregulation by negative feedback from the NTD and identifies a novel gain of function mutant with broad applicability to future Mg^2+^ signaling research.

## INTRODUCTION

Magnesium ions (Mg^2+^) are the most abundant divalent cations in eukaryotes, playing universal roles in myriad cell functions (Jahnen-Dechent & Ketteler, 2012). Within mitochondria, Mg^2+^ is an important protein stabilizing cofactor, forms biologically functional Mg^2+^-ATP complexes and regulates crucial enzymatic activities. Such roles are achieved through two unique properties of Mg^2+^: (i) the ability to form chelates with important intracellular anionic-ligands (*i.e.* small molecule or large biomolecule), and (ii) the capability to compete with calcium ions (Ca^2+^) for binding sites on proteins and membranes (Berridge *et al*, 2000; Chaigne-Delalande *et al*, 2013; Moomaw & Maguire, 2008). The effects of Mg^2+^ on Ca^2+^-handling proteins significantly influence intracellular Ca^2+^ dynamics and signaling (Clapham, 2007; de Baaij *et al*, 2015; Gregan *et al*, 2001).

Total cellular Mg^2+^ concentrations range between ∼17-30 mM; however, concentrations of free Mg^2+^ in the cytosol are estimated between ∼0.5-1.5 mM (Jung *et al*, 1990; Romani, 2011; Rutter *et al*, 1990). Intracellular Mg^2+^ concentrations are strongly buffered and regulated by the combined action of Mg^2+^ binding molecules, Mg^2+^ storage in organelles, and the action of Mg^2+^ channels and exchangers. Remarkably, stored endoplasmic reticulum (ER) Mg^2+^ can be mobilized into the mitochondria through the action ligands such as L-lactate, dramatically modifying metabolism (Daw *et al*, 2020). Mg^2+^ can also alter the electrophysiological properties of ion channels such as voltage-dependent Ca^2+^ channels and potassium (K^+^) channels and affect the binding affinity of Ca^2+^ to EF-hand containing proteins (Glancy & Balaban, 2012; Pilchova *et al*, 2017). All ATP-related biochemical reactions in cells are dependent on Mg^2+^ (Romani, 2007), and extracellular Mg^2+^ also regulates numerous channels such as glutamate receptors and *N*-methyl-D-aspartate (NMDA) receptors (Kumar, 2015). Additionally, this divalent cation contributes to the maintenance of genome stability as a cofactor in DNA repair and protection (Hartwig, 2001). Unsurprisingly, perturbations in intracellular Mg^2+^ concentrations can cause serious cellular dysfunction. For example, decreases in intracellular free Mg^2+^ lead to defective immune responses (Chaigne-Delalande *et al*., 2013; Kanellopoulou *et al*, 2019; Zhou & Clapham, 2009), mutations in the Na^+^/Mg^2+^ exchanger causing chronic intracellular Mg^2+^ deficiency trigger neuronal damage (Kolisek *et al*, 2013) and overexpression of Mg^2+^ channels is a hallmark of several types of cancer (Trapani & Wolf, 2019), to name a few.

Mitochondria have an inner mitochondrial membrane (IMM), separating the mitochondrial matrix (MM) from the intermembrane space (IMS), and an outer mitochondrial membrane (OMM), enclosing the entire organelle. Unregulated, the highly negative IMM potential (∼-180 mV) (Marchi & Pinton, 2014) would drive catastrophically high concentrations of Mg^2+^ entry into the matrix; nevertheless, the matrix Mg^2+^ concentration is similar to the concentration in the cytoplasm, reinforcing that Mg^2+^ influx into the organelle is tightly controlled to maintain optimal mitochondrial function and bioenergetics (Marchi & Pinton, 2014; Pilchova *et al*., 2017).

Residing within the IMM in mammalian cells, <u>m</u>itochondrial <u>R</u>NA <u>s</u>plicing protein <u>2</u> (MRS2) constitutes a major Mg^2+^ entry channel into the MM. Deletion of IMM-localized MRS2 abolishes Mg^2+^ influx into the MM, inducing functional defects in mitochondria and promoting cell death (Merolle *et al*, 2018; Piskacek *et al*, 2009). MRS2 belongs to the large heterogeneous CorA/Mrs2/Alr1 protein superfamily of Mg^2+^ transporters. This family is characterized by the highly conserved Gly-Met-Asn (GMN) motif at the end of the first transmembrane helix, essential for Mg^2+^ transporter function. Mutations of the GMN motif either completely abolish Mg^2+^ transport or profoundly change the ion selectivity of the channel (Knoop *et al*, 2005; Merolle *et al*., 2018; Palombo *et al*, 2013). Human MRS2 contains a large, amino terminal domain (NTD) oriented within the MM, corresponding to residues 58-333 and consisting of ∼71% of the mature polypeptide chain, two transmembrane (TM1 and TM2) domains connected by a highly conserved IMS loop and a smaller, carboxyl terminal domain (CTD) also oriented within the MM (**Fig. 1A**).

**Fig. 1.**
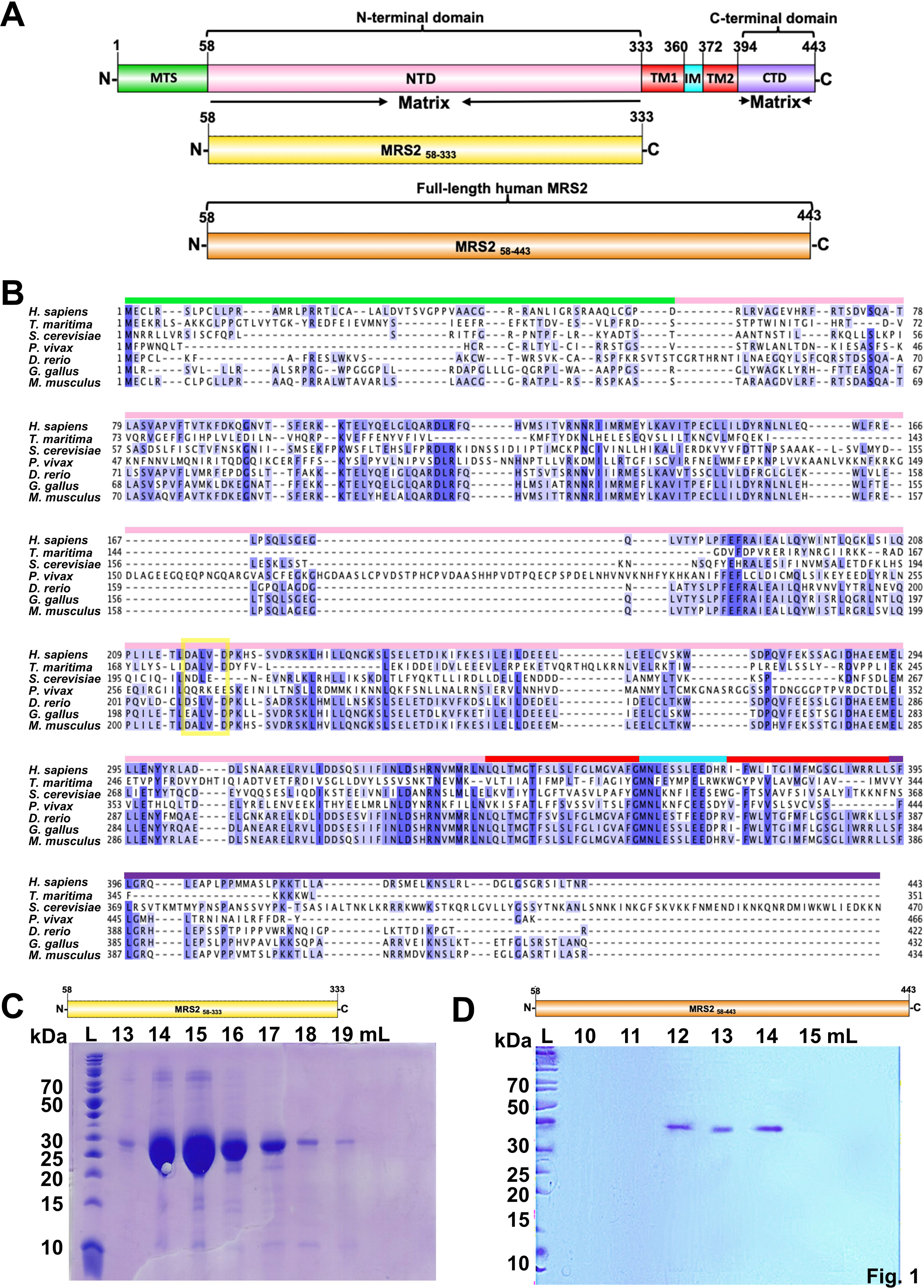
Domain architecture and sequence alignments of MRS2 family proteins. **(A)** Domain architecture of human MRS2. The relative locations of the mitochondrial targeting sequence (MTS, green), amino terminal domain (NTD, magenta), transmembrane 1, 2 (TM1/2, red), intermembrane space (IM, cyan) and C-terminal domain (CTD, violet) are shown. The residue ranges are shown above and below each domain, labeled based on UniProt (Q9HD23) and bioinformatics annotations (see Results). Below, the N-terminal domain (NTD, yellow) and full-length (orange) constructs engineered in the present study are shown relative to the entire pre-protein. **(B)** Multiple sequence alignment of MRS2 orthologues. Sequences for human (NCBI accession Q9WZ3), *Thermotoga maritima* (Q9WZ31), *Saccharomyces cerevisiae* (Q01926), *Plasmodium Vivax* (A0A1G4H438), *Danio rerio (*E7F680), *Gallus gallus* (A0A1D5P665) and *Mus musculus (*Q5NCE8) were aligned in Jalview using T-COFFEE with defaults (Notredame *et al*., 2000; Waterhouse *et al*, 2009). Coloured bars above the human MRS2 sequence mark the boundaries as per the pre-protein in (*A*). The yellow box highlights the location of the DALVD sequence. **(C)** Coomassie blue-stained SDS-PAGE gel showing the elution fractions containing human MRS2_58-333_. **(D)** Coomassie blue-stained SDS-PAGE gel showing the elution fractions containing MRS2_58-443_. In (*C*) and (*D*), elution volumes through an S200 10/300 GL SEC column are shown at top and ladder ‘L’ bands are shown at left.

Although CorA and Mrs2, orthologues of MRS2 in bacteria and yeast, respectively, have been structurally resolved at high resolution (Eshaghi *et al*, 2006; Guskov *et al*, 2012; Johansen *et al*, 2022; Khan *et al*, 2013; Lunin *et al*, 2006; Matthies *et al*, 2016; Payandeh & Pai, 2006; Pfoh *et al*, 2012), low sequence similarity exists between CorA, Mrs2 and MRS2, especially in the NTDs (Zsurka *et al*, 2001). Specifically, the sequence similarity between human MRS2 and yeast (*Saccharomyces cerevisiae*) Mrs2 is 55.4% (and only 20.1% through the NTD), while the sequence similarity between human MRS2 and bacterial (*Thermotoga maritima*) CorA is 43.3% (and only 17.0% through the NTD) (**Fig. 1B**). To reveal how the prominent MRS2 NTD governs the assembly and function of the full channel, we generated recombinant human NTD protein (MRS2_58-333_; residues 58-333) and full-length human MRS2 (MRS2_58-443_; residues 58-443). Using light scattering and chromatographic approaches, we find that in the absence of divalent cations, the NTD self-associates into a homodimer under dilute conditions, while both Mg^2+^ and Ca^2+^, but not cobalt (Co^2+^), suppress the self-association of the domain. In contrast, Co^2+^ disassembles full-length MRS2, while Mg^2+^ and Ca^2+^ have no effect on stoichiometry. 8-Anilino-1-naphthalene sulfonate (ANS) and intrinsic fluorescence measurements suggest that Mg^2+^ and Ca^2+^ bind to distinct sites on the NTD with ∼μM and mM affinity, respectively. Importantly, we identify the D216 and D220 as critical residues for Mg^2+^ coordination to the human MRS2 NTD, where mutating these residues decreases Mg^2+^ affinity ∼7-fold, abrogates Mg^2+^-binding induced increases in α-helicity and solvent accessible hydrophobicity and suppresses the Mg^2+^-induced disassembly of the NTD. Finally, using both permeabilized and intact cell models, we show that Mg^2+^ binding to the NTD suppresses the rate of Mg^2+^ uptake into mitochondria as negative feedback. Collectively, our data reveal previously unknown mechanistic insights underlying human MRS2 autoregulation by the large NTD, which has important implications for understanding the crosstalk between MM Mg^2+^ concentrations, bioenergetics and cell death.

## RESULTS

### MRS2 NTD homodimer assembly is sensitive to divalent cations

Given that human *MRS2* encodes only two putative TMs (**Fig. 1A**) and must oligomerize to form a channel pore, we first evaluated the stoichiometry of the MRS2 NTD (*i.e.* MRS2_58-333_) using size exclusion chromatography with in-line multi-angle light scattering (SEC-MALS). Recombinant MRS2_58-333_ was successfully expressed and isolated with high yield (*i.e.* ∼8 g/L of culture) and purity (**Fig. 1C**). Although the theoretical monomeric molecular weight of MRS2_58-333_ is ∼32.2 kDa, SEC-MALS revealed that, in the absence of divalent cations, MRS2_58-333_ consistently self-associates into a homodimer with estimated molecular weights of 60.9 ± 1.8 kDa and 61.3 ± 0.50 kDa at 2.5 and 5.0 mg/mL, respectively (**Fig. 2A and 2B**). Homodimer formation is apparently tight as single elution peaks and no protein concentration dependence on the molecular weight or elution volume in the 0.45 – 5 mg/mL range was observed (**Table S1**).

**Fig. 2.**
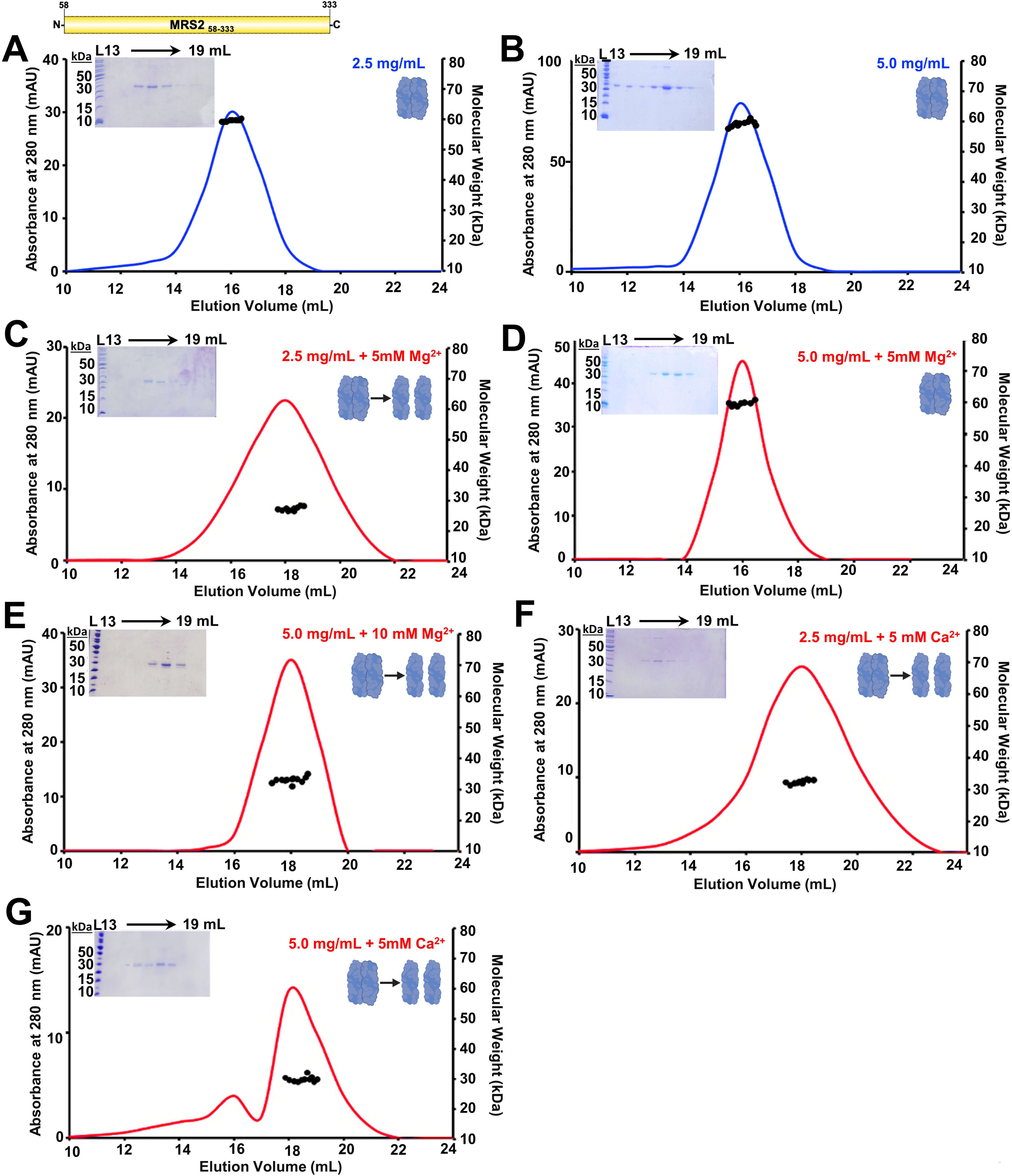
Quaternary structure of MRS2_58-333_ (NTD). **(A, B)** SEC-MALS analysis of MRS2_58-333_ injected at **(A)** 2.5 mg/mL and **(B)** 5.0 mg/mL in the absence of divalent cations. **(C, D)** SEC-MALS data of MRS2_58-333_ injected at **(C)** 2.5 mg/mL and **(D)** 5.0 mg/mL in the presence of 5 mM MgCl_2_. **(E)** SEC-MALS data of MRS2_58-333_ injected at **(E)** 5.0 mg/mL MRS2_58-333_ in the presence of 10 mM MgCl_2_. **(F, G)** SEC-MALS data of MRS2_58-333_ injected at **(F)** 2.5 mg/mL and **(G)** 5.0 mg/mL in the presence of 5 mM CaCl_2_. In (*A-G*), MALS-determined molecular weights are shown through the elution peaks (black circles), left insets show Coomassie-blue stained SDS-PAGE gels of the elution fractions from the 5.0 mg/mL injections, right insets depict the dimerization state of the protein and divalent cation-free and -supplemented chromatograms are coloured blue and red, respectively. Elution volumes are indicated at top and ladder (L) molecular weights at left of the gels. Data are representative of n=3 separate injections from three protein preparations (**Table S1**) and were acquired using an S200 10/300 GL column in 20 mM Tris, 150 mM NaCl, 1 mM DTT, pH 8.0, 10 °C.

SEC-MALS was further used to assess the sensitivity of the homodimer assembly to Mg^2+^ and Ca^2+^, since earlier studies showed divalent cation binding to the CorA NTD regulates channel structure and function (Khan *et al*., 2013; Pfoh *et al*., 2012) (see also Discussion). The presence of 5 mM MgCl_2_ in the elution buffer transitioned the molecular weight of MRS2_58-333_ to monomer at 2.5 mg/mL, with a SEC-MALS molecular weight of 29.4 ± 6.0 kDa (**Fig. 2C**). In contrast, MRS2_58-333_ remained dimeric at 5 mg/mL in the presence of 5 mM MgCl_2_, with a molecular weight of 60.06 ± 0.47 kDa (**Fig. 2D**). Adding 10 mM MgCl_2_ to the 5 mg/mL sample did, however, result in a monomeric molecular weight of 32.6 ± 0.5 kDa (**Fig. 2E**). Remarkably, adding 5 mM CaCl_2_ to both the 2.5 and 5.0 mg/mL MRS2_58-33_ samples robustly caused monomer formation, with measured molecular weights of 31.4 ± 1.0 and 29.0 ± 0.44 kDa, respectively (**Fig. 2F and 2G**). Supplementation with 5 mM MgCl_2_ to MRS2_58-333_ samples at less than 2.5 mg/mL was sufficient to cause monomerization, and shifts to later elution volumes were consistent with all divalent cation-dependent disassembly observations (**Table S1**).

### Divalent cations regulate MRS2 assembly in a domain-specific manner

Since SEC-MALS was performed at 10 °C and is accompanied by a large column dilution (*i.e.* minimum ∼20-fold) that could affect assembly, we used dynamic light scattering (DLS) to assess the distribution of hydrodynamic radii (R_h_) at 1.25 mg/mL in the absence of dilution and at higher temperature (*i.e.* 20 and 37 °C). Bimodal distributions of R_h_ centered at ∼4 and 40 nm were observed for MRS2_58-333_ in the absence of divalent cations at both 20 and 37 °C (**Fig. 3A-3F)**. The addition of either 5 mM CaCl_2_ (**Fig. 3A and 3B**) or 5 mM MgCl_2_ (**Fig. 3C and 3D**) at both temperatures eliminated the larger size distributions. The loss of larger R_h_ was supported qualitatively by earlier decays in the autocorrelation functions when compared to divalent cation-free protein samples (**Fig. 3A-3D, insets**). The bacterial orthologue of MRS2, CorA, can transport Co^2+^ and Mg^2+^, and Co^2+^ is found in trace levels in mammals (Czarnek *et al*, 2015; Guskov & Eshaghi, 2012; Tapiero *et al*, 2003); thus, we also assessed the sensitivity of the MRS2 NTD assembly to Co^2+^. The distribution of R_h_ were not affected by the addition of 5 mM CoCl_2_ at either 20 or 37 °C (**Fig. 3E and 3F**).

**Fig. 3.**
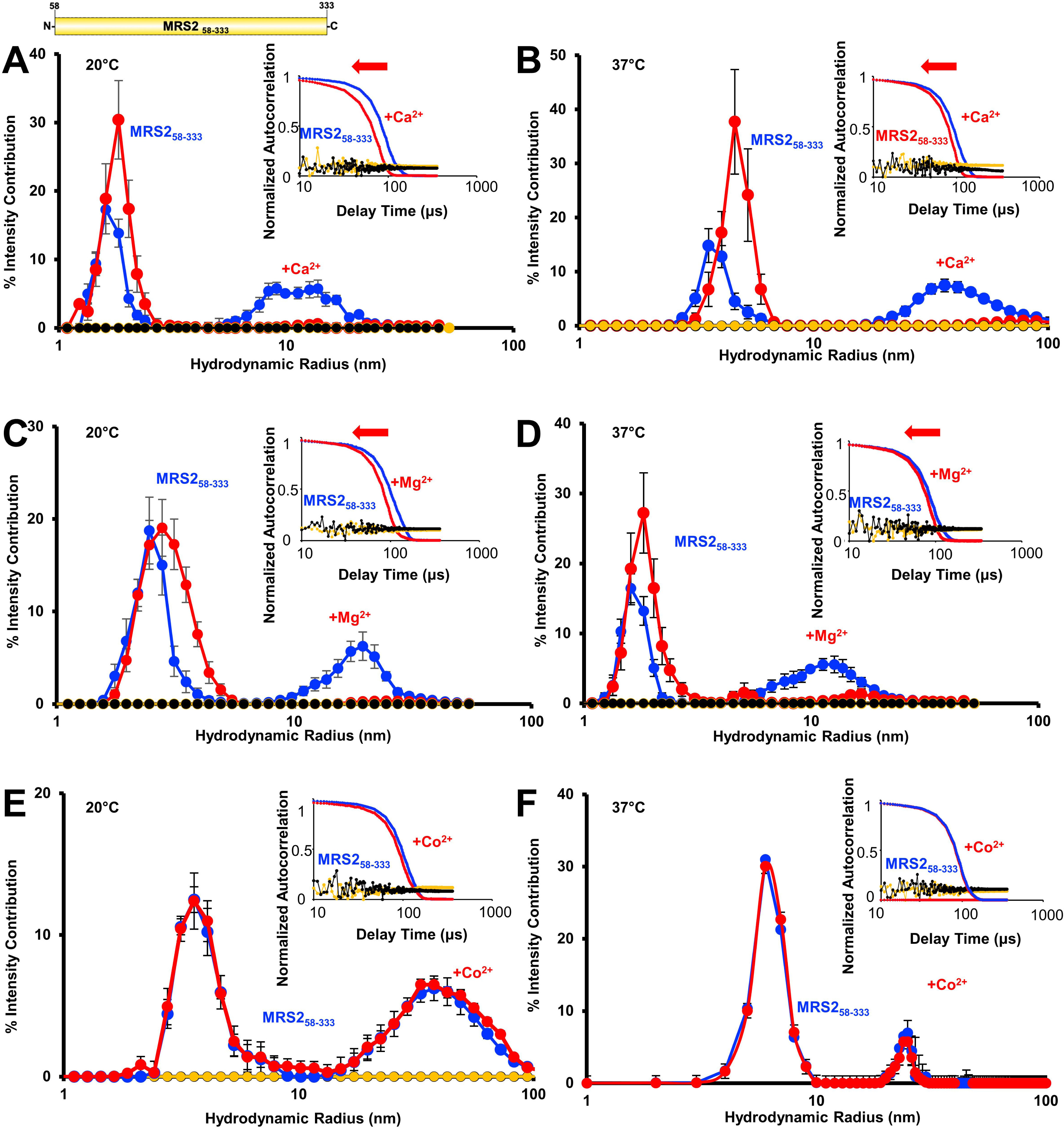
Higher order oligomerization of MRS2_58-333_ (NTD). **(A, B)** The distributions of hydrodynamic radii (R_h_) from the regularization deconvolution of the autocorrelation functions in the presence and absence of 5 mM CaCl_2_ at **(A)** 20 °C and **(B)** 37°C. **(C, D)** The distributions of R_h_ from the regularization deconvolution of the autocorrelation functions in the presence and absence of 5 mM MgCl_2_ at **(C)** 20 °C and **(D)** 37 °C. **(E, F)** The distributions of R_h_ from the regularization deconvolution of the autocorrelation functions in the presence and absence of 5 mM CoCl_2_ at **(E)** 20 °C and **(F)** 37 °C. In (*A-F*), insets show the divalent cation-induced shifts in the autocorrelation functions, divalent cation-free protein sample data are coloured blue, divalent cation-supplemented protein sample data are red, divalent cation-free buffer control data are black and divalent cation-supplemented buffer control data are yellow. Inset data are representative, while deconvoluted R_h_ profiles are means ± SEM of n=3 separate samples from three protein preparations. All data were acquired at 1.25 mg/mL protein in 20 mM Tris, 150 mM NaCl, 1 mM DTT, pH 8.0.

Next, we evaluated the effects of Mg^2+^, Ca^2+^ and Co^2+^ on the assembly of the full-length protein. Full length MRS2, excluding the mitochondrial targeting sequence, (MRS2_58-443_), was successfully expressed and isolated with high purity (**Fig. 1D**). The experimental buffer for MRS2_58-443_ included 3-cholamidopropyl dimethylammonio 1-propanesulfonate (CHAPS) and showed autocorrelation functions consistent with the presence of ∼1-1.5 nm micelles at 37 °C (**Fig. 4A-4C**). MRS2_58-443_ samples showed autocorrelation functions with later decay times compared to buffer alone, which were deconvoluted to R_h_ distributions centered at ∼4 nm and ∼20 nm at 37 °C (**Fig. 4A-4C**). In contrast to the NTD data, changes in autocorrelation functions and size distributions were not observed with MRS2_58-443_ when supplemented with either 5 mM MgCl_2_ or 5 mM CaCl_2_ (**Fig. 4A and 4B)**. Remarkably, 5 mM CoCl_2_ completely abrogated the larger R_h_ distributions, which was qualitatively supported by autocorrelation function shifts to earlier decay times (**Fig. 4C**).

**Fig. 4.**
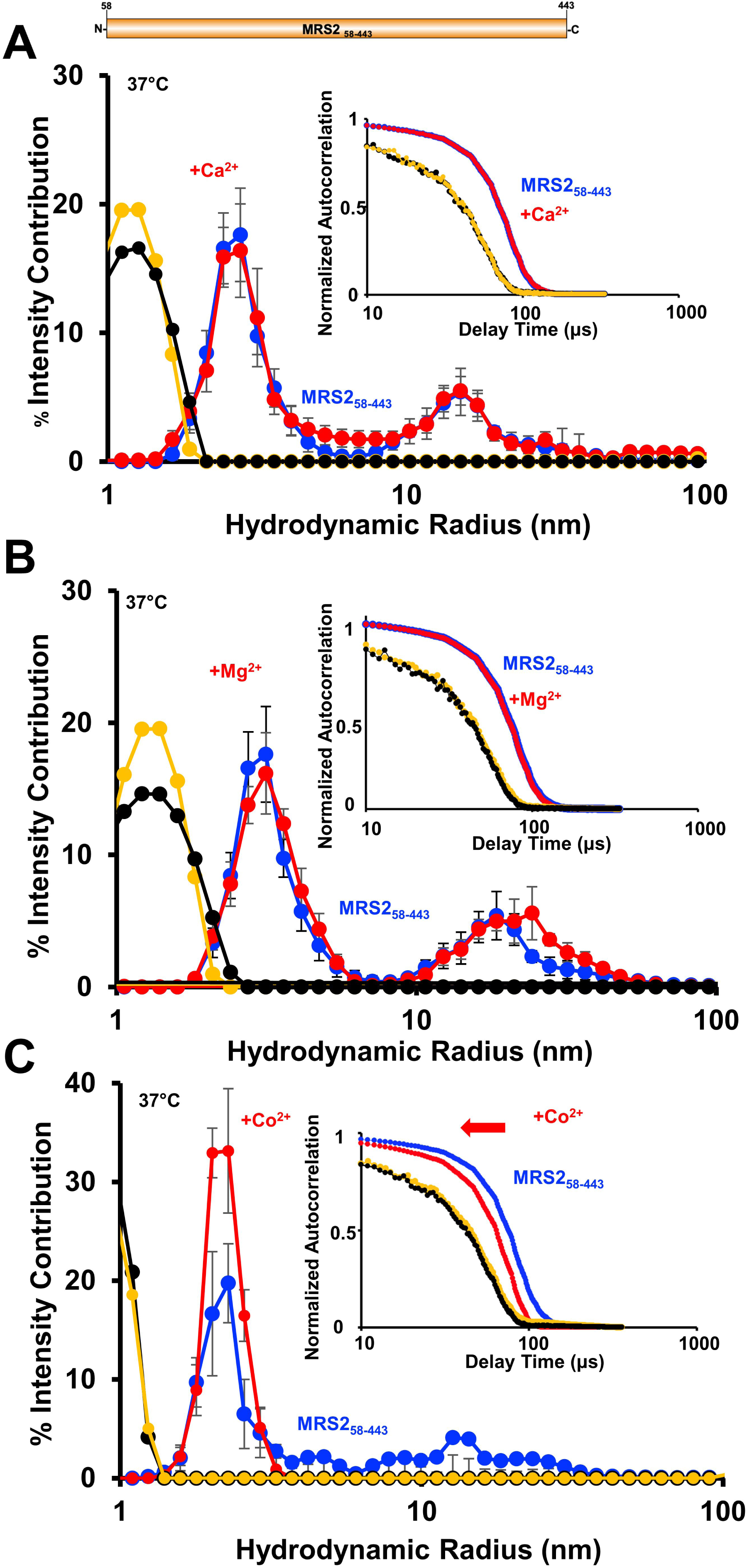
Higher order oligomerization of MRS2_58-443_ (full-length). **(A)** The distributions of R_h_ from the regularization deconvolution of the autocorrelation functions in the presence and absence of 5 mM CaCl_2_. **(B)** The distributions of R_h_ from the regularization deconvolution of the autocorrelation functions in the presence and absence of 5 mM MgCl_2_. **(C)** The distributions of R_h_ from the regularization deconvolution of the autocorrelation functions in the presence and absence of 5 mM CoCl_2_. In (*A-C*), insets show the divalent cation-induced shifts in the autocorrelation functions, divalent cation-free protein sample data are coloured blue, divalent cation-supplemented protein sample data are red, divalent cation-free buffer control data are black and divalent cation-supplemented buffer control data are yellow. Inset data are representative, while deconvoluted R_h_ profiles are means ± SEM of n=3 separate samples from three protein preparations. All data were acquired at 0.5 mg/mL in 20 mM Tris, 150 mM NaCl, 1 mM DTT, 10 mM CHAPS, pH 8.0, 37 °C.

Applying a cumulative deconvolution to extract one weight-averaged R_h_ from all autocorrelation functions reinforced the regularization/polydisperse deconvolution trends described above. Specifically, CoCl_2_ caused a robust decrease in the weight-averaged R_h_ for full-length MRS2, but not the NTD; MgCl_2_ and CaCl_2_ decreased R_h_ for the NTD, but not full-length MRS2 (**Table S2**). Collectively, these data suggest a domain-specific sensitivity to divalent cations, where NTD disassembly is promoted by Mg^2+^ and Ca^2+^, while Co^2+^ de-oligomerizes full length MRS2 due to a sensitivity outside the NTD.

### Mg^2+^ and Ca^2+^ bind to distinct sites on the MRS2-NTD with disparate affinities

Given that both Mg^2+^ and Ca^2+^ dissociate MRS2_58-333_, which contains 3×Trp and 7×Tyr residues, we next used changes in intrinsic fluorescence to evaluate divalent cation binding. Fluorescence emission spectra were acquired using an excitation wavelength of 280 nm as a function of increasing MgCl_2_, CaCl_2_ and CoCl_2_ concentrations. The intensities of the fluorescence emission spectra decreased as a function of increasing MgCl_2_ (**Fig. 5A**) and CaCl_2_ (**Fig. 5B**) concentration. Both MgCl_2_ and CaCl_2_ effects were saturable; however, the intensity decreased by ∼32 % with Mg^2+^ compared to only ∼5 % with Ca^2+^, suggesting distinct structural effects and/or binding sites. In contrast, a titration with CoCl_2_ caused small increases in fluorescence of ∼2 %. Fitting the binding curves to a one-site binding model that accounts for protein concentrations revealed apparent equilibrium dissociation constants (K_d_)s of ∼0.14 ± 0.03, 1.01 ± 0.26 and 0.68 ± 0.30 mM for Mg^2+^, Ca^2+^ and Co^2+^ interactions, respectively (**Table S3**).

**Fig. 5.**
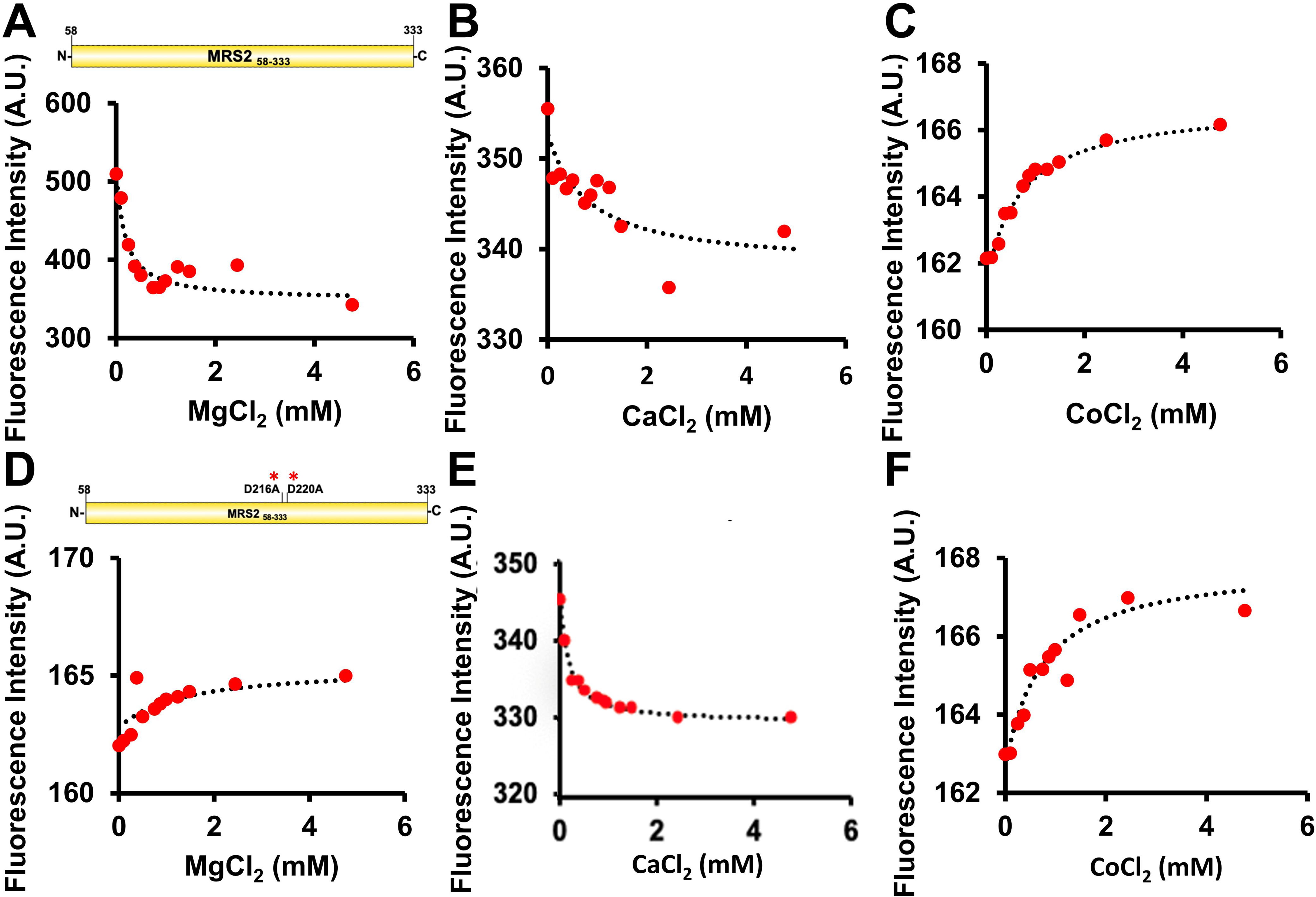
Divalent cation binding affinity to MRS2_58-333_ and MRS2_58-333_ D216A/D220A (NTD). **(A)** Changes in intrinsic fluorescence emission intensity of MRS2_58-333_ as a function of increasing MgCl_2_ concentration. **(B)** Changes in intrinsic fluorescence emission intensity of MRS2_58-333_ as a function of increasing CaCl_2_ concentration. **(C)** Changes in intrinsic fluorescence emission intensity of MRS2_58-333_ as a function of increasing CoCl_2_ concentration. **(D)** Changes in intrinsic fluorescence emission intensity of MRS2_58-333_ D216A/D220A as a function of increasing MgCl_2_ concentration. **(E)** Changes in intrinsic fluorescence emission intensity of MRS2_58-333_ D216A/D220A as a function of increasing CaCl_2_ concentration. **(F)** Changes in intrinsic fluorescence emission intensity of MRS2_58-333_ D216A/D220A as a function of increasing CoCl_2_ concentration. In (*A-F*), data (red circles) show intrinsic fluorescence intensities at 330 nm as a function of increasing divalent cation concentration and are representative of n=3 separate experiments (**Table S3**) performed from three protein preparations. The dashed lines through the data are fits to a one-site binding model that accounts for protein concentration. All experiments were performed with 0.1 μM protein in 20 mM Tris, 150 mM NaCl, 1 mM DTT, pH 8.0 at 22.5 ℃.

We next attempted to pinpoint the residues involved in Mg^2+^ coordination using the CorA crystal structure (4EED.pdb) (Pfoh *et al*., 2012) as a guide. Note that available yeast Mrs2 structures do not resolve any Mg^2+^ ions bound to the NTD. The CorA crystal structure shows that two Asp residues separated by three residues (*i.e.* **<u>D</u>**ALV**<u>D</u>**) are involved in Mg^2+^ coordination at one site. Remarkably, the DALVD sequence is fully conserved in human MRS2 but not yeast (**Fig. 1B**). Thus, after creating a D216A/D220A MRS2_58-333_ double mutant, we reassessed divalent cation binding by intrinsic fluorescence. Not only did the D216A/D220A mutant show a small increase in fluorescence (Mg^2+^ causes a large decrease in the fluorescence intensity of WT MRS2_58-333_; see above), but also a markedly suppressed intensity change as a function of increasing MgCl_2_, consistent with perturbation of Mg^2+^ binding (**Fig. 5D**). In contrast, the CaCl_2_ and CoCl_2_ effects were similar to data acquired using WT MRS2_58-333_ (**Fig. 5E and 5F**). Indeed, fitting the datasets to one-site binding models, revealed apparent equilibrium dissociation constants (K_d_) of ∼0.98 ± 0.25, 0.74 ± 0.49 and 1.37 ± 0.51 mM (**Table S3**), consistent with disruption of the Mg^2+^ interactions but not Ca^2+^ or Co^2+^.

Taken together, these data suggest that Mg^2+^ and Ca^2+^ bind to distinct sites on the MRS2-NTD, with D216 and D220 mediating interactions with Mg^2+^.

### Mg^2+^ enhances while Ca^2+^ suppresses solvent exposed hydrophobicity of MRS2 NTD

Given the changes in stoichiometry observed by DLS and SEC-MALS, we next assessed the solvent exposed hydrophobicity of MRS2_58-333_ in the absence and presence of Mg^2+^ and Ca^2+^ by monitoring extrinsic ANS fluorescence. ANS binds to solvent accessible hydrophobic regions on biomolecules, resulting in a blue shifted fluorescence emission maximum and increased intensity (Stryer, 1965). Baseline fluorescence emission spectra of ANS in the presence of buffer alone were insensitive to the addition of 5 mM MgCl_2_ or 5 mM CaCl_2_ (**Fig. S1A and S1B**). Indeed, ANS binding was detected in the presence of 2.5 mg/mL MRS2_58-333_, as evidenced by the blue shifted fluorescence emission maximum and increased intensity compared to the buffer controls (**Fig. S1A and S1B**). Supplementing the protein samples with 5 mM MgCl_2_ caused a small but significant increase in ANS fluorescence intensity, suggesting enhanced exposed hydrophobicity (**Fig. 6A and 6B**). Conversely, supplementation with 5mM CaCl_2_ caused a small but significant decrease in ANS fluorescence intensity, indicating decreased solvent exposed hydrophobicity (**Fig. 6C and 6D**).

**Fig. 6.**
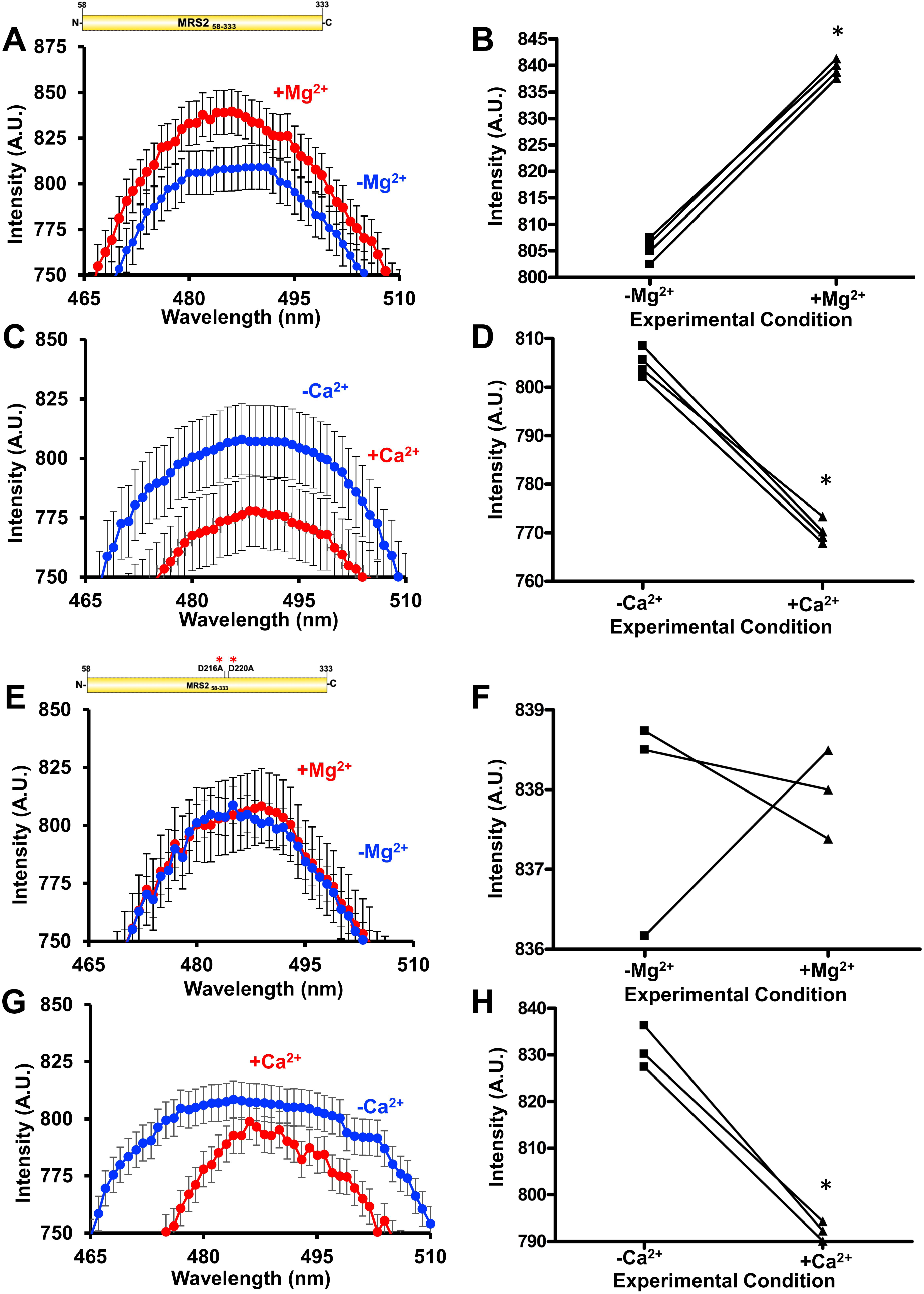
Solvent-exposed hydrophobicity of MRS2_58-333_ and MRS2_58-333_ D216A/D220A (NTD). **(A, B)** ANS fluorescence emission spectra of **(A)** MRS2_58-333_ in the absence (blue) and presence (red) of 5 mM MgCl_2_ and **(B)** paired statistical comparisons of the peak intensities. **(C, D)** ANS fluorescence emission spectra of **(C)** MRS2_58-333_ in the absence (blue) and presence (red) of 5 mM CaCl_2_ and **(D)** paired statistical comparisons of the peak intensities. **(E, F)** ANS fluorescence emission spectra of **(E)** MRS2_58-333_ D216A/D220A in the absence (blue) and presence (red) of 5 mM MgCl_2_ and **(F)** paired statistical comparisons of the peak intensities. **(G, H)** ANS fluorescence emission spectra of **(G)** _MRS258-333_ D216A/D220A in the absence (blue) and presence (red) of 5 mM CaCl_2_ and **(H)** paired statistical comparisons of the peak intensities. In (*A, C, E* and *G*), data are means ± SEM of n=3 separate samples from three protein preparations. In (*B, D, F* and *H*), comparisons are paired Student’s t-test analyses, where * *p* < 0.05. ANS binding experiments were performed using 30 μM protein and 50 μM ANS in 20 mM Tris, 150 mM NaCl, 1 mM DTT, pH 8.0 at 15 °C.

We next performed a similar set of experiments with the D216A/D220A MRS2_58-333_ protein. Consistent with our observation that this double mutant disrupts Mg^2+^ but not Ca^2+^ binding to the NTD, ANS emission spectra in the presence of protein showed no differences with or without MgCl_2_ supplementation (**Fig. 6E and 6F**; **Fig. S1C**), while CaCl_2_ supplementation caused a small but significant decrease in ANS fluorescence intensity (**Fig. 6G and 6H**; **Fig. S1D**). This ANS binding data reinforces the notion of disparate Ca^2+^ and Mg^2+^ binding sites and suggests that these divalent cations may cause distinct MRS2 NTD conformational changes.

### D216A/D220A mitigates Mg^2+^-dependent disassembly of the MRS2 NTD

Next, we tested whether the D216A/D220A double mutation could abolish the Mg^2+^-dependent monomerization and decreased R_h_ observed with WT MRS2 NTD. In the absence of the divalent cation, SEC-MALS revealed that D216A/D220A MRS2_58-333_ elutes as a homodimer with a molecular weight of 59.7 ± 2.0 kDa when injected at 2.5 mg/mL (**Fig. 7A**), similar to WT MRS2_58-333_ (**Table S1**). In contrast to WT evaluated at 2.5 mg/mL, however, the SEC-MALS-determined molecular weight of the double mutant remained dimeric (*i.e.* 59.0 ± 2.4 kDa) after the addition 5 mM MgCl_2_ (**Fig. 7B**). We also assessed whether Mg^2+^ could alter R_h_ of the D216A/D220A MRS2_58-333_ by DLS. Addition of 5 mM MgCl_2_ neither altered the distribution of R_h_ nor the autocorrelation function compared to samples evaluated in the absence of the cation (**Fig. 7C**). Note that a bimodal distribution of R_h_ centered at ∼4 nm and ∼40 nm was observed with the D216A/D220A MRS2_58-333_ protein (**Fig. 7C**), similar to WT.

**Fig. 7.**
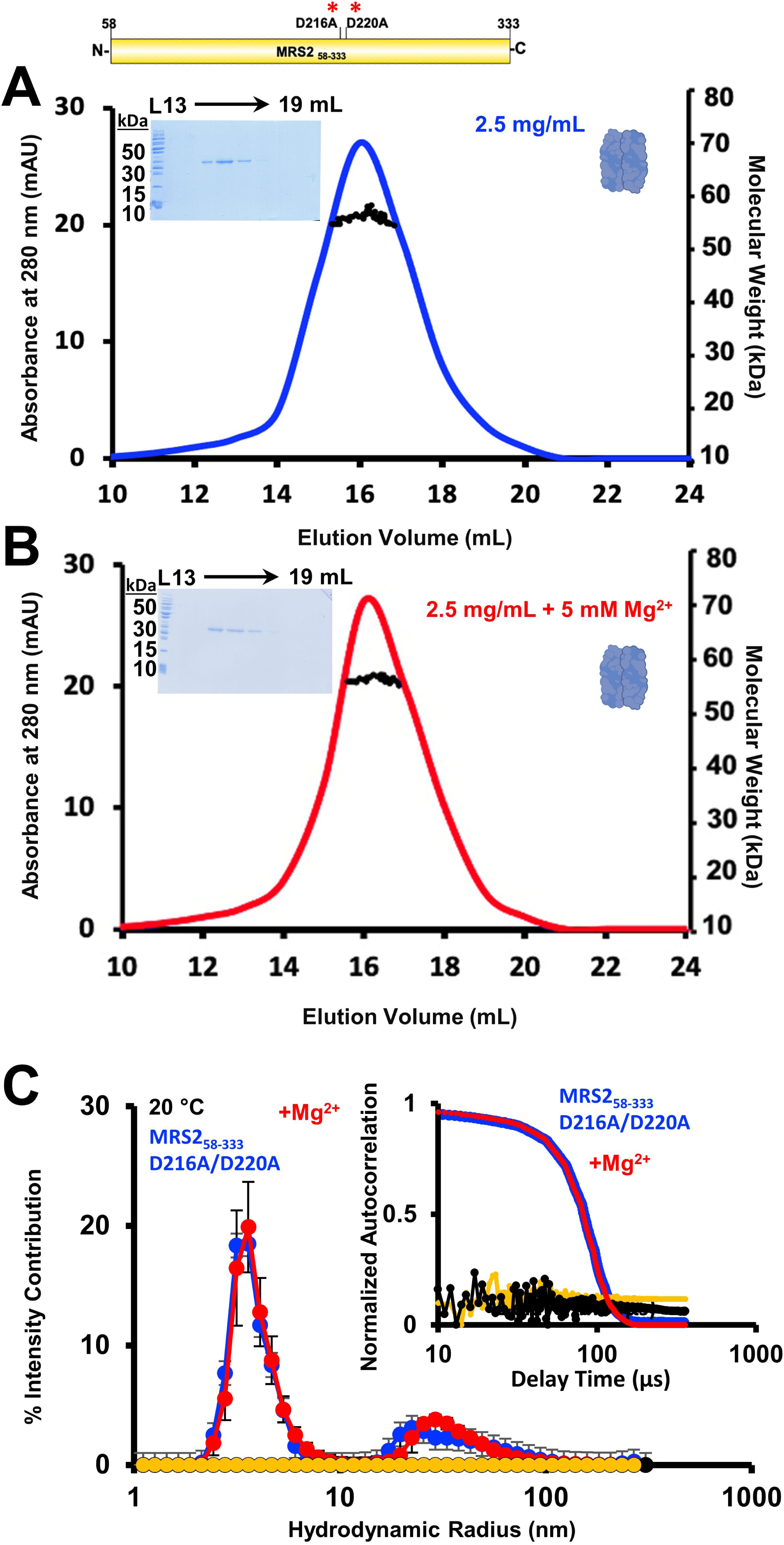
Quaternary structure and higher order oligomerization of MRS2_58-333_ D216A/D220A (NTD). **(A, B)** SEC-MALS analysis of MRS2_58-333_ D216A/D220A injected at 2.5 mg/mL in the **(A)** absence or **(B)** presence of 5 mM MgCl_2_. **(C)** DLS analysis of MRS2_58-333_ D216A/D220A at 1.25 mg/mL. The distributions of R_h_ from the regularization deconvolution of the autocorrelation functions are shown in the presence and absence of 5 mM MgCl_2_ at 20 °C. In (*A-B*), MALS-determined molecular weights are shown through the elution peaks (black circles), left insets show Coomassie-blue stained SDS-PAGE gels of the elution fractions from the 2.5 mg/mL injections and right insets depict the dimerization state of the protein. Elution volumes are indicated at top and ladder ‘L’ molecular weights at left of the gels. Data are representative of n=3 separate injections from three protein preparations (**Table S1**) and were acquired using an S200 10/300 GL column in 20 mM Tris, 150 mM NaCl, 1 mM DTT, pH 8.0, 10 °C. In (*C*), inset show the Mg^2+^-induced shift in the autocorrelation functions, Mg^2+^-free protein sample data are coloured blue, Mg^2+^-supplemented protein sample data are red, Mg^2+^-free buffer control data are black and Mg^2+^-supplemented buffer control data are yellow. Inset data are representative, while deconvoluted R_h_ profiles are means ± SEM of n=3 separate samples from three protein preparations. Data were acquired in 20 mM Tris, 150 mM NaCl, 1 mM DTT, pH 8.0, 20 °C.

Together, these light scattering analyses demonstrate that Mg^2+^-dependent disassembly of the MRS2 NTD requires the D216 and D220 residues, where double mutation to Ala abrogates quaternary structure sensitivity to the cation.

### D216A/D220A abrogates increased α-helicity and thermal stability in the MRS2 NTD caused by Mg^2+^ binding

Having observed that Mg^2+^ binding affects quaternary and tertiary levels of MRS2-NTD structure, we next used far-UV circular dichroism (CD) spectroscopy to assess secondary structure. At 37°C, MRS2_58-333_ displayed well-defined mean residue ellipticity (MRE) minima at ∼208 and ∼222 nm, indicating high levels of α-helicity (**Fig. 8A**). Remarkably, addition of 5 mM MgCl_2_ directly to the cuvette resulted in an increase in α-helicity, evidenced by more intense negative ellipticity at ∼208 and 222 nm (**Fig. 8A and 8C**). Similar results were observed for MRS2_58-333_ at 20 °C (**Fig. S2A**).

**Fig. 8.**
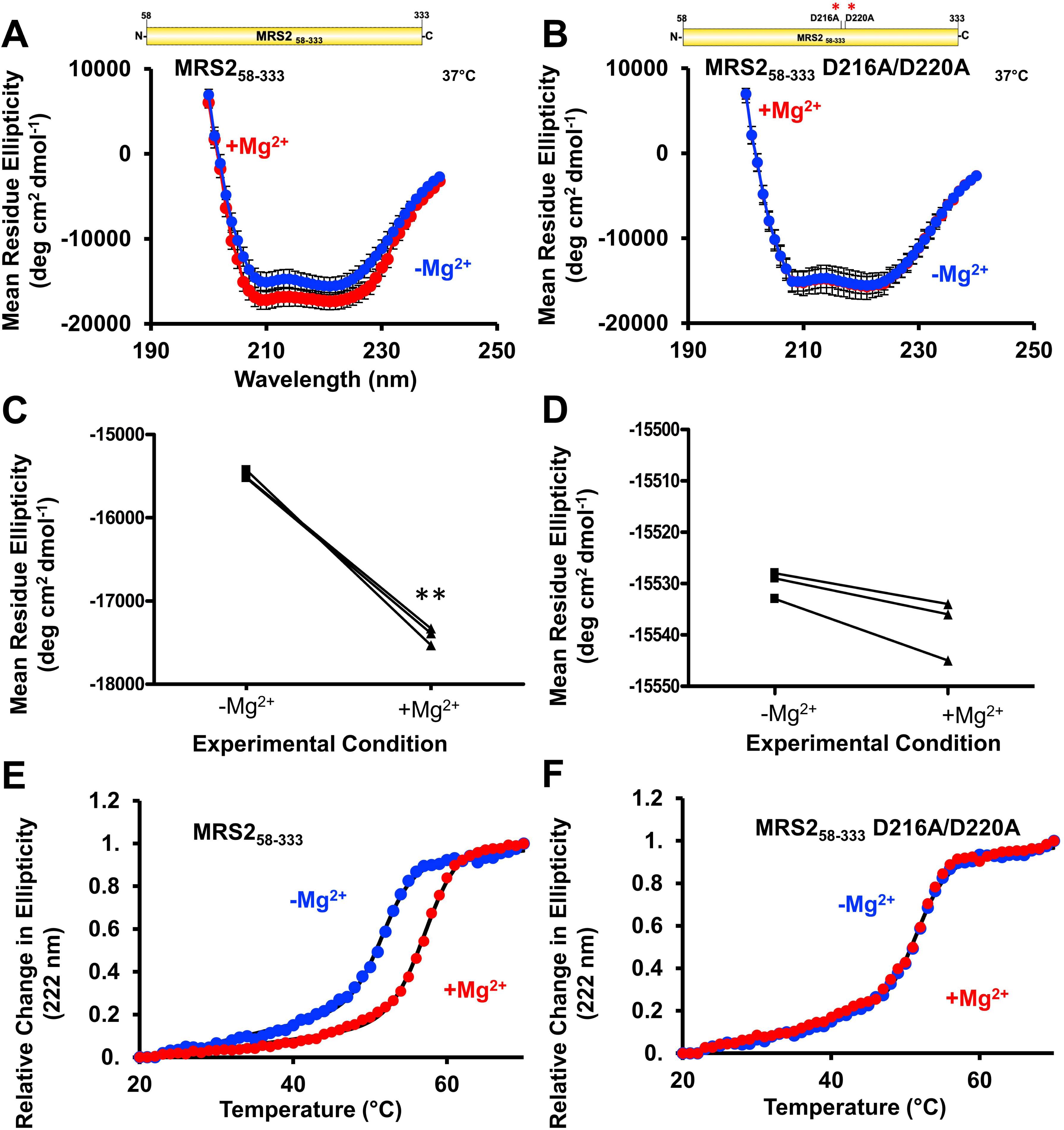
Secondary structure and thermal stability of MRS2_58-333_ and MRS2_58-333_ D216A/D220A (NTD). **(A, B)** Far-UV CD spectra of **(A)** MRS2_58-333_ and **(B)** MRS2_58-333_ D216A/D220A in the absence (blue) and presence (red) of 5 mM MgCl_2_. **(C, D)** Statistical comparisons of Mg^2+^-induced changes in mean residue ellipticity (MRE) at 222 nm for **(C)** MRS2_58-333_ and **(D)** MRS2_58-333_ D216A/D220A. **(E, F)** Changes in MRE (222 nm) as a function of increasing temperature (*i.e.* thermal stability) in the absence (blue) and presence (red) of 5 mM MgCl_2_ for **€** MRS2_58-333_ and **(F)** MRS2_58-333_ D216A/D220A. In (*A-B*), data are means ± SEM of n=3 experiments with samples from three protein preparations. In (*C-D*), comparisons are paired Student’s t-test analyses of data from (*A* and *B*), where ** *p* < 0.01. In (*E-F*), data (circles) are representative of n=3 separate experiments with samples from three protein preparations, and solid black lines are Boltzmann sigmoidal fits through the data to extract apparent midpoints of temperature denaturation (T_m_). All far-UV CD experiments were acquired with 0.5 mg/mL protein in 20 mM Tris, 150 mM NaCl, 1 mM DTT, pH 8.0, with spectra acquired at 37 °C.

To gain further evidence that D216 and D220 play a critical role in Mg^2+^ binding to the NTD, we also acquired far-UV CD spectra using D216A/D220A MRS2_58-333_. The far-UV CD spectrum of the double mutant showed a similar level of negative ellipticity as WT with two well-defined minima at ∼208 and ∼222 nm (**Fig. 8A and 8B**), suggesting that secondary structure folding was not perturbed by the D216A/D220A substitutions. Unlike WT, adding 5 mM MgCl_2_ directly to the cuvette did not significantly alter the ellipticity for the double mutant (**Fig. 8D**). Unchanging spectra after 5 mM MgCl_2_ addition were also observed for the double mutant at 20 °C (**Fig. S2B**).

We next evaluated thermal stability by monitoring the change in far-UV CD ellipticity at 222 nm as a function of increasing temperature. The thermal melts of MRS2_58-333_ acquired in the absence of Mg^2+^ exhibited a mean Boltzmann sigmoidal fitted midpoint of temperature denaturation (T_m_) of 51 ± 0.62 °C (**Fig. 8E**). Protein samples supplemented with 5 mM MgCl_2_ were stabilized by ∼7 °C as the mean T_m_ shifted to 58 ± 0.36 °C (**Fig. 8E**). Thermal melt experiments with the D216A/D220A MRS2_58-333_ protein revealed similar mean T_m_ values of 52 ± 0.70 °C and 52 ± 0.64 °C in the presence and absence of Mg^2+^, respectively (**Fig. 8F**).

Collectively, these data reveal that Mg^2+^ binding stabilizes the MRS2 NTD, consistent with an observed increase in α-helicity. Further, the structural and stability augmentation is dependent on D216 and D220 as mutation of these residues renders the NTD insensitive to Mg^2+^, reinforcing the importance of these sites to coordinating Mg^2+^.

### Mg^2+^ binding to the MRS2 NTD negatively regulates mitochondrial Mg^2+^ uptake

To link our *in vitro* observations with MRS2 function, we monitored Mg^2+^ dynamics using Mag-Green in HeLa cells overexpressing WT and D216A/D220A MRS2. HeLa cells were incubated with the membrane permeant Mag-Green-AM to cytosolically load the cells with the Mg^2+^ sensitive dye. After washing and bathing the cells with intracellular buffer (IB), the plasma membrane (PM) was permeabilized with 5 μM digitonin, and 3 mM MgCl_2_ was added to the bath. Mitochondrial Mg^2+^ uptake rates were inferred from the clearance of cytosolic Mg^2+^, measured as the decrease in Mag-Green fluorescence, as has been previously done (Daw *et al*., 2020). At 20 °C and after MgCl_2_ addback, digitonin-permeabilized HeLa cells transfected with empty pCMV vector (control), pBSD-MRS2 (WT) and pBSD-MRS2 D216A/D220A (mutant), all showed increases in Mag-Green fluorescence followed by a decay associated with Mg^2+^ clearance (**Fig. S3A**). Fitting the data to a single exponential decay indicated greater cytosolic Mg^2+^ clearance rates for cells expressing the mutant MRS2 compared to WT MRS2 expressing or control cells (**Fig. S3B and S3C**).

We performed a similar set of Mg^2+^ clearance assays at 37 °C since past work demonstrated greater MRS2 activity at this temperature (Daw *et al*., 2020). Digitonin-permeabilized control, WT- and mutant-MRS2 expressing HeLa cells all showed increases in Mag-Green fluorescence after 3 mM MgCl_2_ addback, followed by a decay associated with Mg^2+^ clearance from the cytosol (**Fig. S3D**). Fitting the data to a single exponential decay indicated greater cytosolic Mg^2+^ clearance rates for WT MRS2-expressing cells compared to control cells and mutant MRS2-expressing cells compared to control and WT MRS2-expressing cells (**Fig. S3E and S3F**).

Collectively, these data suggest that MRS2 overexpression enhances cytosolic Mg^2+^ clearance, and Mg^2+^ interactions with the MRS2 NTD act as a negative feedback switch to temper Mg^2+^ uptake into the mitochondria.

### Gain of function D216K/D220K mutant relieves negative feedback on MRS2 activity

To probe whether mutation of the Mg^2+^ binding site causes a *bona fide* gain of function, we reconstituted human WT and D216K/D220K MRS2 in WT and Mrs2 knockout (Mrs2^-/-^) hepatocytes. Primary murine hepatocytes were transfected with empty vector, human MRS2-mRFP or MRS2 D216K/D220K-mRFP plasmids. Twenty-four hours post-transfection, a genetically-encoded, mitochondrially targeted Mag-FRET biosensor (*i.e.* mito-Mag-FRET) was transduced into the cells to directly measure mitochondrial Mg^2+^ uptake. Confocal images of the transfected/transduced WT hepatocytes show strong co-expression and co-localization of MRS2 and MRS2 D216K/D220K with the cerulean and citrine fluorescence of the mito-Mag-FRET biosensor, indicating mitochondrial localization of the human WT and mutant MRS2 in murine cells (**Fig. 9A**). Human WT and mutant MRS2 were similarly well-expressed and strongly co-localized with the mito-Mag-FRET fluorophores in the Mrs2^-/-^ hepatocytes (**Fig. 9B**). As expected, a 10 mM MgCl_2_ bolus resulted in an increased mito-Mag-FRET signal in WT cells (**Fig. 9C**). While WT hepatocytes expressing human MRS2 showed a similar mito-Mag-FRET response to the MgCl_2_ bolus, cells transfected with human MRS2 D216K/D220K exhibited highly potentiated mitochondrial Mg^2+^ uptake compared to controls (**Fig. 9D**). Next, we evaluated the mito-Mag-FRET responses in Mrs2^-/-^ hepatocytes. Human MRS2 was fully capable of functionally reconstituting the Mg^2+^ channel in Mrs2^-/-^ hepatocyte mitochondria. Remarkably, the MRS2 D216K/D220K formed channels that greatly enhanced mitochondrial Mg^2+^ uptake compared to WT human MRS2 (**Fig. 9E**). Given the striking potentiation of Mg^2+^ uptake in hepatocytes co-expressing the D216K/D220K mutant but not WT human MRS2 with endogenous Mrs2, our data suggest that the Mg^2+^ binding-deficient MRS2 mutant dominantly mediates a gain of mitochondrial Mg^2+^ uptake function.

**Fig. 9.**
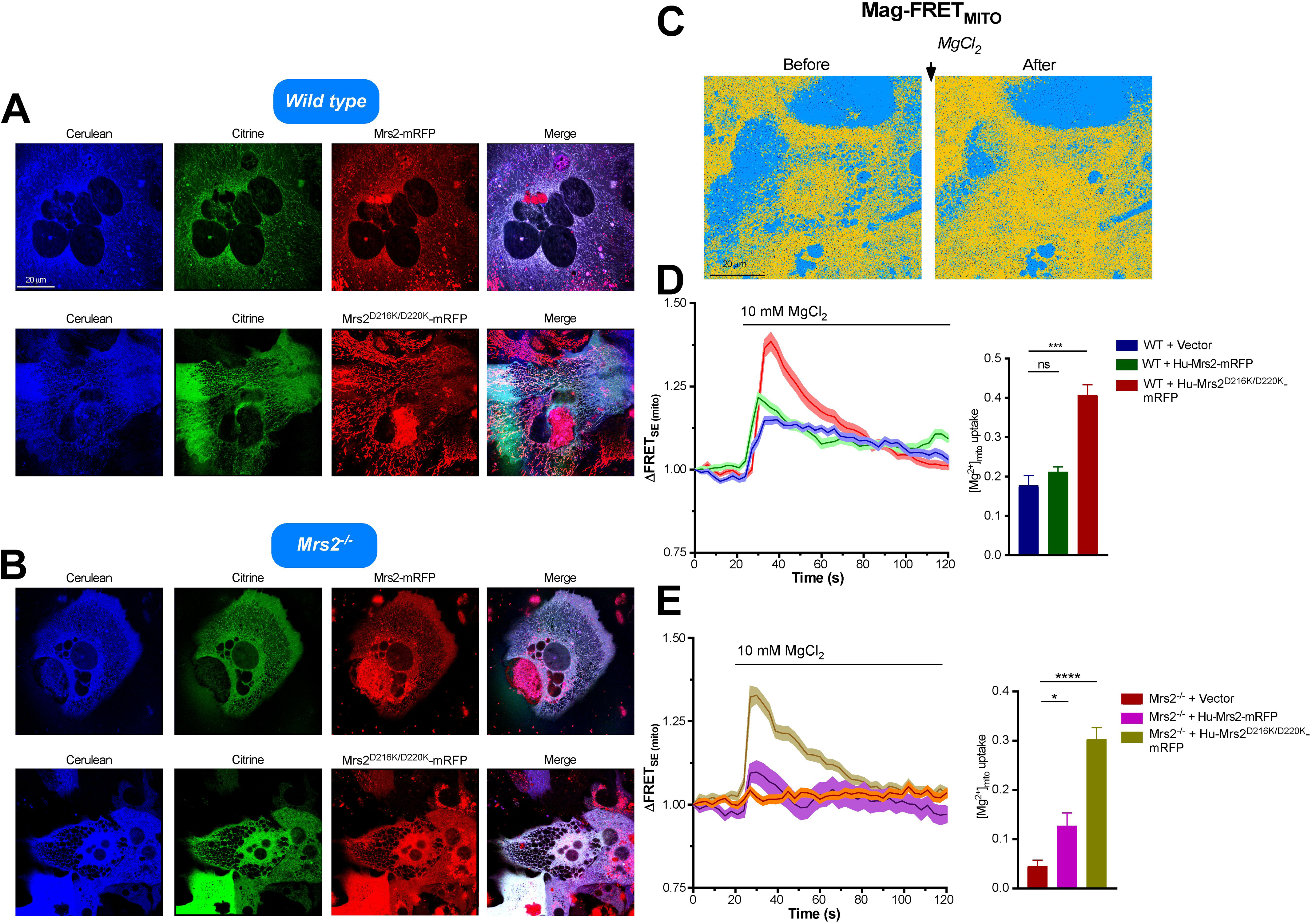
The D216K/D220K double mutant enhances human MRS2 channel activity. **(A, B)** Representative confocal images showing expression and mitochondrial localization of human WT MRS2-mRFP and MRS2 D216K/D220K-mRFP in WT **(A)** or Mrs2^-/-^ **(B)** primary murine hepatocytes transduced with the adenoviral mito-Mag-FRET sensor. **(C)** Representative FRET images before and after 10 mM MgCl_2_ addition to primary hepatocytes expressing mito-Mag-FRET. **(D)** Mean traces (left panel) showing relative mito-Mag-FRET ratio changes upon addition of 10 mM MgCl_2_ and comparisons of peak mitochondrial Mg^2+^ uptake responses in WT hepatocytes (right panel). **(E)** Mean traces (left panel) showing relative mito-Mag-FRET ratio changes upon addition of 10 mM MgCl_2_ and comparisons of peak mitochondrial Mg^2+^ uptake responses in Mrs2^-/-^ hepatocytes (right panel). In (*A-E*), all measurements were performed at 37°C and quantified values are normalized as FRET/FRET_0_, where FRET and FRET_0_ are the signals at any time and initial timepoints, respectively. Unpaired Student’s t-test were performed for comparison of peak responses, where \**p* < 0.05, \*\*\**p* < 0.001, \*\*\*\**p* < 0.0001, ns, not significant. Data are means ± SEM of n=3 for WT + vector, n=5 for WT + Hu-MRS2-mRFP, n=3 for WT + Hu-MRS2^D216K/D220K^-mRFP, n=3 for Mrs2^-/-^ + vector, n=4 for Mrs2^-/-^ + Hu-MRS2-mRFP and n=3 for Mrs2^-/-^ + Hu-MRS2^D216K/D220K^-mRFP, where n is the number of separate transfections.

## DISCUSSION

Human MRS2 belongs to the heterogeneous CorA/Mrs2/Alr1 superfamily of Mg^2+^ transporters, where CorA, Alr1 and Mrs2/MRS2 comprise the principal Mg^2+^ uptake systems in bacteria, yeast plasma membrane and mitochondria, respectively. Bacterial CorA has been the most extensively studied family member, yielding mechanistic and functional insights on these channels (Franken *et al*, 2022; Jin *et al*, 2022). Nevertheless, given the low sequence similarity between human MRS2 and these homologues, there remains a major knowledge gap concerning the precise structural, functional, and regulatory mechanisms of human MRS2. Here, we isolated and biophysically characterized the largest domain of human MRS2, corresponding to the matrix-oriented NTD. We found that MRS2 NTD forms a homodimer under dilute conditions, which may be a building block to higher order oligomers. Remarkably, Mg^2+^ and Ca^2+^ disassembled both higher order MRS2 NTD oligomers and homodimers but not full length MRS2 assemblies. In contrast, Co^2+^ disassembled full length MRS2 oligomers but not MRS2 NTD. We estimated the K_d_ of Mg^2+^ binding to be ∼0.14 mM, and a D216A/D220A MRS2 NTD double mutant disrupted this Mg^2+^ binding but had no effect on Ca^2+^ binding, indicating disparate binding sites for these two divalent cations. Remarkably, this D216A/D220A double mutant abrogated the enhanced solvent exposed hydrophobicity, α-helicity and thermal stability mediated by Mg^2+^ binding. Further, MRS2 NTD oligomers and homodimers harboring this double mutation remained intact in the presence of Mg^2+^. Finally, we showed that reconstitution of D216A/D220A or D216K/D220K MRS2 mutants in mammalian cells greatly increased mitochondrial Mg^2+^ uptake compared to WT MRS2 expressing cells.

Several CorA crystal and cryoelectron microscopy (cryoEM) structures have been elucidated in the presence of divalent cations, revealing a pentameric assembly (Cleverley *et al*, 2015; Eshaghi *et al*., 2006; Guskov *et al*., 2012; Johansen *et al*., 2022; Matthies *et al*., 2016; Nordin *et al*, 2013; Payandeh & Pai, 2006; Pfoh *et al*., 2012). The first TM, which lines the channel pore, and second TM, position the intervening GMN motif for ion binding and selectivity at the pore entrance (Pfoh *et al*., 2012). Upstream of TM1, a large intracellular domain of CorA, analogous to the matrix-oriented human MRS2 NTD, fans out into the cytoplasm and is composed of eight α-helices and a six-stranded β-sheet (*Thermotoga maritima*; 4EED.pdb) (**Fig. 10A**). For *T. maritima* CorA, two Mg^2+^ binding sites (M1 and M2) have been identified per intracellular domain (Pfoh *et al*., 2012). M1 is made up of D89 and D253, while M2 is comprised of D175 and D179. Whereas earlier studies indicated that a symmetrizing of the pentameric intracellular domain assembly upon Mg^2+^ binding to the NTD closes the channel (Matthies *et al*., 2016; Pfoh *et al*., 2012), more recent work indicates both symmetric and asymmetric assemblies are formed in the presence and absence of Mg^2+^, and channel conductance is dependent on lowered symmetric state population and coupled with a reduced energy barrier to an ensemble of open states in low Mg^2+^ (Johansen *et al*., 2022; Kowatz & Maguire, 2019). Nevertheless, it is evident that Mg^2+^ binding increases the rigidity/decreases the dynamics of the CorA intracellular domain (Chakrabarti *et al*, 2010; Johansen *et al*., 2022; Pfoh *et al*., 2012; Rangl *et al*, 2019).

**Fig. 10.**
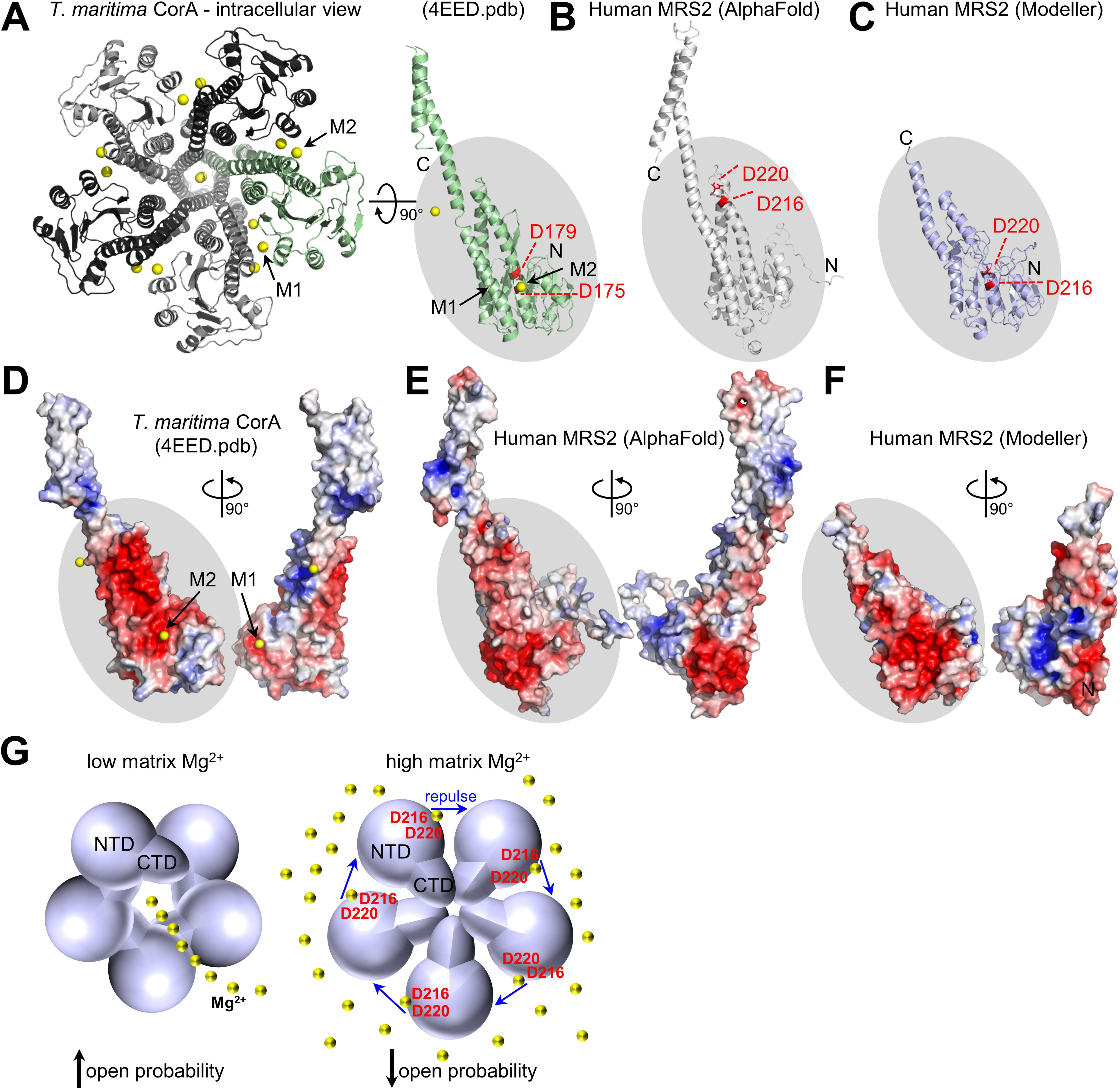
Model of Mg^2+^-induced negative feedback in MRS2 function. **(A)** At left, intracellular view of the experimentally determined *T. maritima* CorA pentamer structure (4EED.pdb). A backbone cartoon representation of each protomer is shown with M1 and M2 Mg^2+^ binding sites indicated (black arrows). At right, view of a single protomer (green), rotated 90° relative to the pentamer view. Positions of D175 and D179, equivalent to human D216 and D220, are coloured red and labeled (broken lines). **(B)** AlphaFold model of a human MRS2 protomer, with the D216 and D220 positions coloured red and labeled (broken lines). **(C)** Modeller-derived model of the human MRS2 NTD, using the experimentally determined CorA structure (4EED.pdb) as the template. The D216 and D220 positions are coloured red and labeled (broken lines). **(D, E, F)** Electrostatic surface potential maps of the **(D)** CorA, **(E)** AlphaFold MRS2 and **(F)** human MRS2 NTD protomers. The blue to red color gradient represents a +5 kT/e to -5 kT/e potential gradient, respectively. (G) Model of matrix Mg^2+^-induced inhibition of human MRS2 open probability. At low matrix Mg^2+^ (left), the human MRS2 channel has a high open probability; at high matrix Mg^2+^ (right), Mg^2+^ binding to a site mediated by D216 and D220 repulses NTD interactions, leading to a low open probability. Despite weak NTD interactions at high Mg^2+^, the channel remains assembled via TM and/or CTD interactions. A pentameric human MRS2 channel is depicted in homology to CorA, but the functional stoichiometry of the human MRS2 channel remains unknown. In (*B-F*), the same relative view is highlighted by the grey oval, Mg^2+^ ions are represented by yellow spheres, the amino and carboxyl termini are labeled N and C, respectively, and Mg^2+^ binding sites are indicated as M1 and M2.

The CorA M1 Mg^2+^ binding residues do not appear conserved in human MRS2 based on EMBOSS pairwise and T-COFFEE multiple sequence alignment (**Fig. 1B**) (Notredame *et al*, 2000; Olson, 2002). The AlphaFold (Jumper *et al*, 2021) model of human MRS2 places the M2 binding residues closer to the PM region of the NTD, inconsistent with the near β-sheet position elucidated in the CorA crystal and cryoEM structures (**Fig. 10B**). Thus, we created a homology model in Modeller (Webb & Sali, 2017), positioning the DALVD residues, conserved with CorA, in a structurally similar position as elucidated by the CorA crystal structures (**Fig. 10C**). Our model suggests M2 coordinates Mg^2+^ near the interface between subunits, analogous to CorA; however, whereas CorA E88 and D89 creates an acidic surface on the opposite face of the subunit, complementing basic Mg^2+^ from two sides (**Fig. 10D**), our model shows a more basic surface, which could be repulsed by Mg^2+^ and cause subunit disassembly (**Fig. 10E and 10F**). Note that the sequence identity between human MRS2 NTD and the *T. maritima* CorA intracellular domain is low (*i.e.* ∼17 %), and so a high-resolution structure of human MRS2 is needed to reveal the precise mechanism for Mg^2+^-induced disassembly characterized in the present study.

Mg^2+^ increased α-helicity and stability of the human MRS2 NTD, consistent with past NMR data showing decreased backbone dynamics of CorA in the presence of high Mg^2+^ (Johansen *et al*., 2022). The far-UV CD spectra reported here resemble previous data from our lab, where we found no effect by MgCl_2_, likely due to variability in protein concentration measurements (Daw *et al*., 2020). Here, we applied MgCl_2_ addback to the same sample to expose the secondary structure change. Several lines of evidence suggest distinct Ca^2+^ and Mg^2+^ binding sites on the MRS2 NTD. First, the change in intrinsic fluorescence caused by the two cations was different; second, whereas Mg^2+^ increased, Ca^2+^ decreased solvent accessible hydrophobicity; third, D216A/D220A double mutant increased the Mg^2+^ K_d_ ∼7-fold, while having no effect on the Ca^2+^ K_d_; finally, the Mg^2+^-dependent solvent accessible hydrophobicity change was abrogated, while the Ca^2+^ response was maintained by the D216A/D220A double mutant. Given the mitochondrial matrix has a free Mg^2+^ concentration of ∼0.5-1.5 mM (Jung *et al*., 1990; Rutter *et al*., 1990), the Mg^2+^ K_d_ of ∼0.14 mM reported here would suggest the MRS2 structure, stability and oligomerization would be sensitive to physiologically relevant fluctuations in Mg^2+^ levels within the matrix. In contrast, Ca^2+^ concentrations would rarely approach mM levels in the matrix, and thus the MRS2 NTD would have to be positioned close to a Ca^2+^ channel pore to be affected (Bauer, 2001; Chad & Eckert, 1984; Tadross *et al*, 2013).

Interestingly, while Co^2+^ dissociated larger full length MRS2 assemblies, we observed no effect on MRS2 NTD by DLS. In contrast, Mg^2+^ and Ca^2+^ did not alter the assembly of full-length MRS2 but dissociated the MRS2 NTD. We posit that MRS2_58-333_ (NTD) within MRS2 full-length undergoes disassembly in the presence of Mg^2+^ and Ca^2+^, while the C-terminal domain and/or TM regions remain interacting. Such a change would be undetectable by DLS, as the complex size would be unaffected. Further, we believe Co^2+^-mediated disassembly occurs via binding to a region outside the NTD. Since estimates for the mitochondrial concentration of Co^2+^ range from ∼50 to 90 nM (Czarnek *et al*., 2015; Tapiero *et al*., 2003), the precise physiological significance of Co^2+^ interactions with any human MRS2 domain remains unclear.

Using permeabilized and intact cells, our data show that Mg^2+^ binding to the MRS2 NTD negatively regulates the channel. Permeabilized cells overexpressing the D216A/D220A double mutant MRS2 cleared Mg^2+^ from the cytosol at increased rates compared to WT MRS2-expressing cells. Further, human WT and D216K/D220K MRS2 were fully capable of reconstituting functional MRS2 channels in intact primary murine Mrs2^-/-^ hepatocytes, with the double mutant causing highly potentiated Mg^2+^ uptake in Mrs2^-/-^ and WT mitochondria, indicative of gain of function activity. Interestingly, a study using CorA harboring mutations aimed at disrupting Mg^2+^ binding to M1 showed WT-like ^63^Ni^2+^ transport (Kowatz & Maguire, 2019). Here, we focused on M2 residues, since M1 does not appear to be conserved in human MRS2, discovering a robust, dominantly increased mitochondrial Mg^2+^ uptake upon disruption of Mg^2+^ binding to the NTD.

In conclusion, our work reveals the large NTD functions as a negative feedback regulator of human MRS2 channel function. We propose Mg^2+^ binding to the NTD, contributed to by D216 and D220, disrupts inter-subunit interactions by negating electrostatically complementary interactions between neighbouring subunits (**Fig. 10G**). Mg^2+^ binding to the MRS2 NTD increases α-helicity, stability and solvent exposed hydrophobicity, which we believe are biophysical changes that propagate to the pore and/or crucial gating residues, resulting in inhibition without disassembly of the channel (**Fig. 10G**). These data distinguish human MRS2 from bacterial CorA observations, where Mg^2+^ binding to the analogous intracellular domain shields electrostatically repulsive interfaces, promoting and bridging a symmetric interaction between intracellular domains (Matthies *et al*., 2016; Pfoh *et al*., 2012).

## MATERIALS AND METHODS

### MRS2 expression and purification

The human MRS2 NTD was identified as residues 58-333 using bioinformatic identification of the mitochondrial targeting sequence (MTS) (Almagro Armenteros *et al*, 2019; Buchan & Jones, 2019; Fukasawa *et al*, 2015), and TM1 and TM2 (Krogh *et al*, 2001), followed by comparison of these predictions with the annotations in UniProt (Accession #Q9HD23). Human MRS2_58-333_ was subcloned out of the BDS vector into pET-28a (Novagen) using PCR and NdeI and XhoI restriction sites. Overnight protein expression at 37 °C from the pET-28a-MRS2_58-333_ vector was done using BL21 (DE3) *Escherichia coli* cells cultured in Luria broth (LB), induced with 0.4 mM isopropyl β-d-1-thiogalactopyranoside (IPTG). Protein was purified under native conditions using HisPur (Thermo-Fisher) nickel-nitrilotriacetic acid beads as per the manufacturer guidelines. The wash and elution buffers contained 20 mM Tris (pH 8.0), 150 mM NaCl, 1 mM DTT and 20 mM Tris (pH 8.0), 150 mM NaCl,1 mM DTT, 300 mM imidazole, respectively. After dialysis in 20 mM Tris (pH 8.0), 150 mM NaCl, 1 mM DTT buffer using a 3,500 Da MWCO membrane (Thermo Scientific), the N-terminal hexa-histidine tag was cleaved with ∼2 Units of bovine thrombin (Sigma) per 1 mg of protein. A final size exclusion chromatography (SEC) step through an S200 10/300 GL column (Cytiva), achieved > 95 % protein purity as assessed by sodium dodecyl sulfate polyacrylamide gel electrophoresis (SDS-PAGE) and Coomassie blue staining.

D216A and D220A mutant were introduced into MRS2_58-333_ by PCR-mediated site-directed mutagenesis and expression and purification for this construct were performed as described for WT-MRS2_58-333_. The complementary mutagenic primers were 5’-CCTTGAGACCTTGGCTGCTTTGGTGGCCCCCAAACATTCTTC-3’ and 3’-GAAGAATGTTTGGGGGCCACCAAAGCAGCCAAGGTCTCAAGG -5’.

Full-length human MRS2 taken as residues 58-443 (MRS2_58-443_) was cloned and expressed using the same approach described for MRS2_58-333_. Purification was performed as described for the NTD, except with the addition of 10 mM CHAPS to both the elution and SEC buffers.

### Size exclusion chromatography with in-line multi angle light scattering

SEC-MALS was performed using a Superdex 200 Increase 10/300 GL column (Cytiva) connected to an AKTA Pure FPLC system (Cytiva). A DAWN HELEOS II detector (Wyatt) and an Optilab Trex differential refractometer (Wyatt) were used to estimate the molecular weight of MRS2_58-333_ under various experimental conditions. The entire in-line FPLC/MALS system was housed in cold cabinet maintained at ∼10 °C. Data were obtained for four different protein concentrations: 0.45 mg/mL, 0.90 mg/mL, 2.5 mg/mL and 5 mg/mL in 20 mM TRIS (pH 8), 150 mM NaCl, and 1 mM DTT, using 100 μL injections of sample at each concentration. MALS molecular weights were determined in the accompanying ASTRA software (version 7.1.4; Wyatt) using Zimm plot analysis and a protein refractive index increment (dn/dc) = 0.185 L/g. Divalent cation containing experiments were performed by supplementing the running buffers and protein samples with 5 or 10 mM MgCl_2_ and CaCl_2_, as indicated.

### Dynamic light scattering

DLS measurements were made with a DynaPro Nanostar (Wyatt) instrument using a scattering angle of 90 °. After centrifugation at 15,000 ×g for 5 min at 4°C, 5 μL of supernatant was loaded into a JC-501 microcuvette and measurements were collected as 10 consecutive acquisition scans with each acquisition being an average of 5 s. MRS2_58-333_ protein samples were assessed at 1.25 mg/mL in 20 mM TRIS (pH 8), 150 mM NaCl, and 1 mM DTT in the absence or presence of 5 mM MgCl_2_, CaCl_2_ or CoCl_2_. Similarly, MRS2_58-443_ protein samples were assessed at 0.5 mg/mL in 20 mM TRIS (pH 8), 150 mM NaCl, 10 mM CHAPS and 1 mM DTT, in the absence or presence of 5 mM MgCl_2_, CaCl_2_ or CoCl_2._ For both proteins, data were acquired at 20 and 37 °C, as indicated. All autocorrelation functions were deconvoluted using the regularization algorithm to extract the polydisperse distribution of hydrodynamic radii (R_h_) and cumulants fit for monodisperse weight-averaged R_h_ using the accompanying DYNAMICS software (version 7.8.1.3; Wyatt).

### Intrinsic fluorescence measurements for cation binding

A Cary Eclipse spectrofluorimeter (Agilent/Varian) was used to acquire intrinsic fluorescence emission spectra. Spectra were acquired for 0.1 mg/mL MRS2_58-333_ in 20 mM TRIS (pH 8), 150 mM NaCl, and 1 mM DTT, using a 600 μL quartz cuvette. The fluorescence emission intensities were recorded at 22.5 °C from 300 to 450 nm, using a 1 nm data pitch and an excitation wavelength of 280 nm. Excitation and emission slit widths were set to 5 and 10 nm, respectively, and the photomultiplier tube (PMT) detector was set to 650 V. Emission spectra were obtained before and after supplementation with increasing concentrations of CaCl_2_, MgCl_2_ or CoCl_2_, added directly to the cuvette. A total of 15 emission spectra were acquired with increasing concentrations of divalent cation between 0 and 5 mM. Spectral intensities at 330 nm were corrected for the dilution associated with the volume change upon each addition to the cuvette, and resultant curves were fit to a one site binding model that takes into account protein concentration using R (version 4.2.1) to extract apparent equilibrium dissociation constants (K_d_).

### Extrinsic 8-anilinonapthalene-1-sulfonic acid fluorescence

Extrinsic ANS fluorescence measurements were performed using a Cary Eclipse spectrofluorometer (Agilent/Varian). Spectra were acquired at 15 °C for 30 μM MRS2_58-333_ in 20 mM TRIS (pH 8), 150 mM NaCl, 1 mM DTT and 0.05 mM ANS, using a 600 μL quartz cuvette. The excitation wavelength was set to 372 nm, and the extrinsic ANS fluorescence emission spectra were acquired from 400 to 600 nm, with the photomultiplier tube (PMT) detector set at 750V. Excitation and emission slit widths were set to 10 and 20 nm, respectively, for all ANS experiments. To monitor divalent cation-induced changes in exposed hydrophobicity of MRS2_58-333_, 5 mM CaCl_2_ or 5 mM MgCl_2_ was added directly into the cuvette. Negligible effects of these cations on free ANS fluorescence were confirmed by acquiring similar spectra in the absence of protein.

### Far-UV Circular dichroism spectroscopy

Far-UV CD spectra were acquired using a Jasco J-810 CD spectrometer with electronic Peltier temperature regulator (Jasco). Each spectrum was taken as an average of 3 accumulations, recorded at 37°C using a 1 mm pathlength quartz cuvette in 1 nm increments, 8 s averaging time and 1 nm bandwidth. To eliminate technical variability in magnitude signals, after acquiring divalent cation free spectra, 5 mM MgCl_2_ was added to the same samples and spectra were re-acquired.

Thermal melts were recorded using a 1 mm pathlength quartz cuvette by monitoring the change in CD signal at 222 nm from 20 – 95 °C. A scan rate of 1°C min^−1^, 1 nm bandwidth and 8 s averaging time was used during data acquisition. Mg^2+^-free and Mg^2+^-supplemented data were fit using a Boltzmann sigmoidal equation to estimate the midpoint of temperature denaturation (T_m_) using R (version 4.2.1).

### Mitochondrial Mg^2+^ uptake experiments using Mag-Green

HeLa cells were cultured in Dulbecco’s modified Eagle’s media (DMEM) with high glucose (Wisent), 10 % (v/v) fetal bovine serum (Sigma), 100 µg/mL penicillin and 100 Units/mL streptomycin (Wisent) at 37 °C in a 5 % CO_2_, 95 % (v/v) air mixture. Cells cultured in 35 mm dishes were transfected with PolyJet transfection reagent (SignaGen) according to the manufacturer guidelines. Following ∼12 h, cells were incubated with 0.725 μM of the Mg^2+^ indicator Mag-Green for 30 minutes at 37°C. Cells were subsequently washed in divalent cation-free phosphate buffered saline (PBS), pH 7.4 and suspended in 2 mL of intracellular buffer (IB) composed of 20 mM HEPES (pH 7), 130 mM KCl, 2 mM KH_2_PO_4_, 10 mM NaCl, 5 mM succinate, 5 mM malate and 1 mM pyruvate. A 20 % (v/v) cell suspension in IB was created in a quartz cuvette. Cytosolic Mag-Green fluorescence was monitored using a PTI QuantMaster spectrofluorimeter (Horiba) equipped with electronic temperature control using excitation and emission wavelengths of 506 and 531 nm, respectively, and excitation and emission slit widths of 2.5 and 2.5 nm, respectively. After a 30 s Mag-Green baseline fluorescence measurement, 2 mM EDTA plus 5 μM digitonin was added to permeabilize the PM. After 300 s, 3 mM MgCl_2_ was added to the cuvette and the Mag-Green signal was measured for 600 s. Mitochondrial Mg^2+^ uptake was correlated with the clearance of Mg^2+^, taken as the decrease in Mag-Green fluorescence after re-introduction of Mg^2+^ into the system, as previously done (Daw *et al*., 2020). The rates of Mag-Green fluorescence decrease were extracted by fitting the traces after the Mg^2+^ addback to a single exponential decay in R (version 4.2.1).

### Mito-Mag-FRET measurements in primary murine hepatocytes

Primary murine hepatocytes isolated from WT and Mrs2^-/-^ (Daw *et al*., 2020) mice grown on 25-mm collagen-coated glass coverslips were transfected with empty vector, human MRS2-mRFP (Hu-MRS2-mRFP) or Hu-MRS2 D216K/D220K-mRFP plasmids. Twenty-four h post-transfection, hepatocytes were transduced with adenoviral mito-Mag-FRET (Daw *et al*., 2020) (20 MOI) for an additional 48 hours. The sub-cellular localization of ectopically expressing MRS2 and Mito-Mag-FRET were visualized using Leica SP8 confocal microscope (Manheim, Germany). The cells were excited using the 405 nm laser line, and the emission was collected using the hybrid detector (HyD). The cerulean channel, 460-490 nm, and citrine channel, 510-550 nm, served to detect the emissions from the fluorescence resonance energy transfer (FRET). FRET emissions were acquired following donor and acceptor excitation sequences. Selected Region of interests (ROIs) were drawn, and the acquired sequences were background corrected for acceptor cross excitation cross-talk, acceptor cross excitation, and FRET cross-talk (α = A/C; γ = B/C; δ = A/B). Time-lapse imaging was performed using the above-described acquisition mode, and the corresponding FRET efficiencies were analyzed. Selective ROIs focused on mitochondrial-targeted mito-Mag-FRET sensor signals, and the captures FRET sensitized emissions (relative FRET_SE_) were plotted.

### Structure modeling and visualization

Ten homology models of the human MRS2 NTD were generated using the experimentally determined CorA structure (4EED.pdb) as the template in MODELLER (version 9.16) (Webb & Sali, 2017). The model with the lowest DOPE score was taken for visualization and structural analyses. Electrostatic surface potential maps were generated using the PDB2PQR (Dolinsky *et al*, 2007) and APBS (Jurrus *et al*, 2018) plugins in PyMOL. Monovalent ionic strength was set to 150 mM and temperature to 310 K for the electrostatic calculation. All structure images were generated using PyMOL (Version 2.4, Schrödinger, LLC).

### Statistics

Unpaired Student’s t-test was used when comparing two independent groups, paired Student’s t-test was used when comparing outcomes of the same group before and after treatment, and one-way ANOVA followed by Tukey’s post-hoc test was used for multiple means comparisons between three or more groups. All non-linear regression fitting and statistical analyses were done in GraphPad Prism (4.03) or R (4.2.1).

## DATA AVAILABILITY

Data available upon request.

## SUPPORTING INFORMATION

This article contains supporting/extended view information, including Table S1, Table S2, Table S3, Figure S1, Figure S2 and Figure S3.

## CRediT AUTHOR STATEMENT

Sukanthathulse Uthayabalan: conceptualization, methodology, formal analysis, investigation, writing – original draft, visualization. Neelanjan Vishnu: Conceptualization, methodology, formal analysis, investigation, writing – original draft, visualization. Muniswamy Madesh: conceptualization, validation, writing – review and editing, resources, supervision, funding acquisition. Peter B. Stathopulos: conceptualization, validation, writing – review and editing, resources, supervision, project administration, funding acquisition.

## ACKNOWLEDGEMENTS

This work was supported by a Canadian Institutes of Health Research Project Scheme Grant 438225 (to PBS) and National Institutes of Health (R01GM109882, R01HL086699, R01HL142673, R01GM135760) and DOD/DHP-CDMRP PR181598P-1 grants (to MM).

## CONFLICT OF INTEREST

The authors declare that they have no conflicts of interest with the contents of this article.

## FIGURE LEGENDS

**Supporting Fig. S1.**
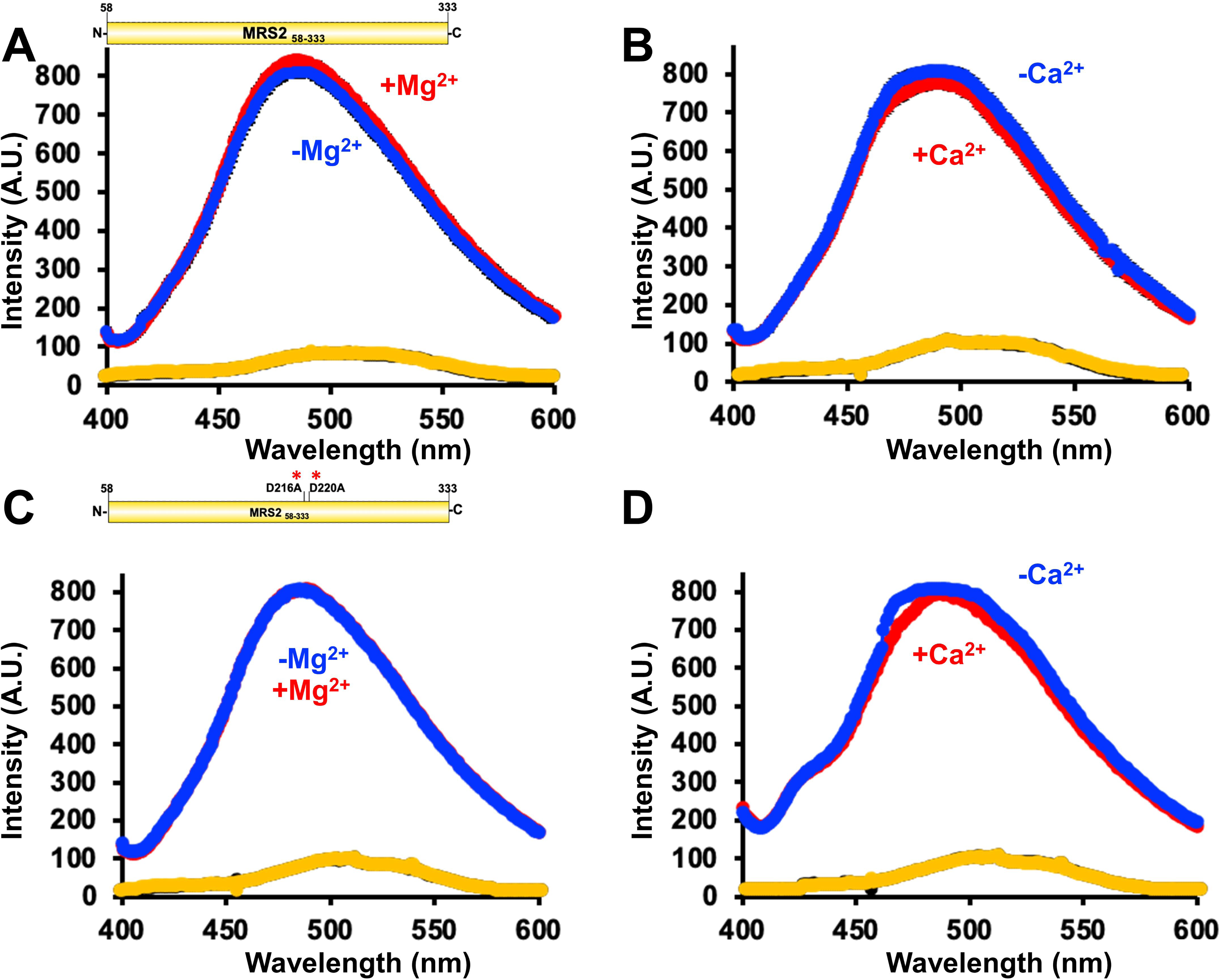
Complete ANS fluorescence emission spectra of MRS258-33 and MRS258-333 D216A/D220A (NTD). **(A)** ANS emission spectra of MRS2_58-333_ in the absence (blue) and presence (red) of 5 mM MgCl_2_. **(B)** ANS emission spectra of MRS2_58-333_ in the absence (blue) and presence (red) of 5 mM CaCl_2_. **(C)** ANS emission spectra of MRS2_58-333_ D216A/D220A in the absence (blue) and presence (red) of 5 mM MgCl_2_. **(D)** ANS emission spectra of MRS2_58-333_ D216A/D220A in the absence (blue) and presence (red) of 5 mM CaCl_2_. In (*A-D*), buffer-only spectra acquired in the absence and presence divalent cation are coloured black and yellow, respectively. Data are means ± SEM of n=3 separate experiments from three protein preparations. Data were acquired using 30 μM protein and 50 μM ANS in 20 mM Tris, 150 mM NaCl, 1 mM DTT, pH 8.0 at 15 °C.

**Supporting Fig. S2.**
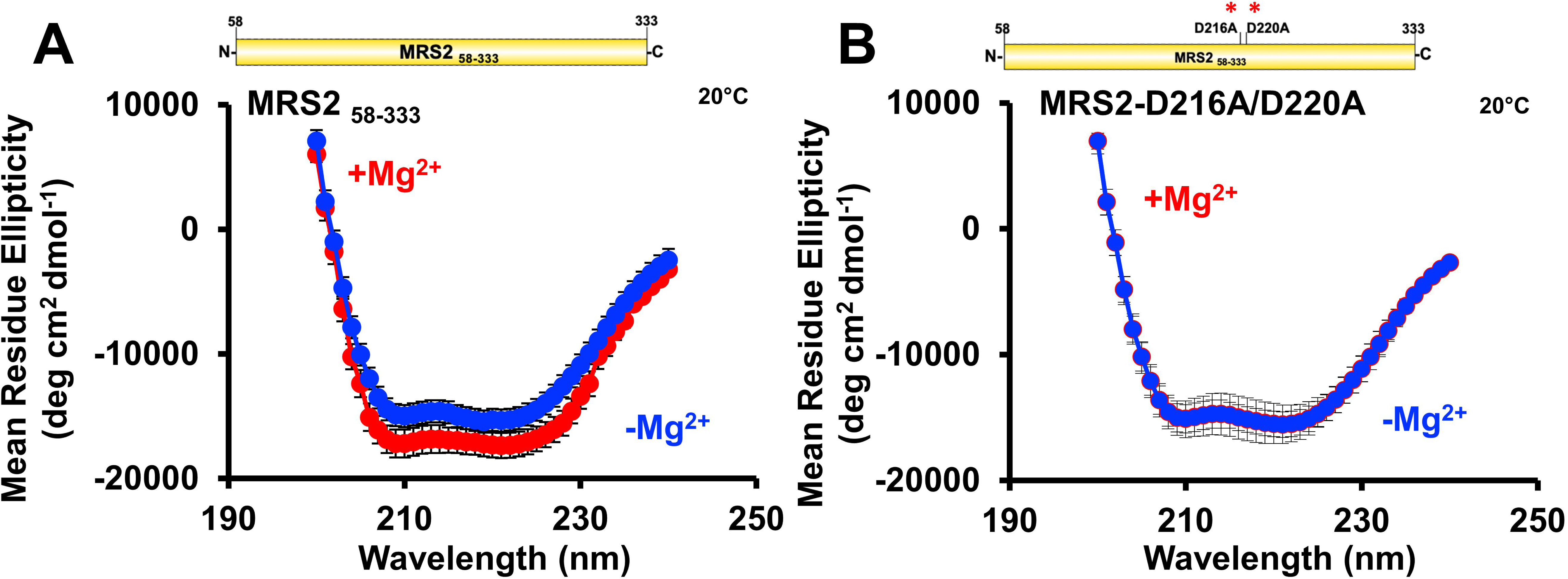
Secondary structure of MRS2_58-333_ and MRS2_58-333_ D216A/D220A (NTD). **(A, B)** Far-UV CD spectra of **(A)** MRS2_58-333_ and **(B)** MRS2_58-333_ D216A/D220A in the absence (blue) and presence (red) of 5 mM MgCl_2_. In (*A-B*), data are means ± SEM of n=3 experiments with samples from three protein preparations. Far-UV CD spectra were acquired with 0.5 mg/mL protein in 20 mM Tris, 150 mM NaCl, 1 mM DTT, pH 8.0, at 20 °C.

**Supporting Fig. S3.**
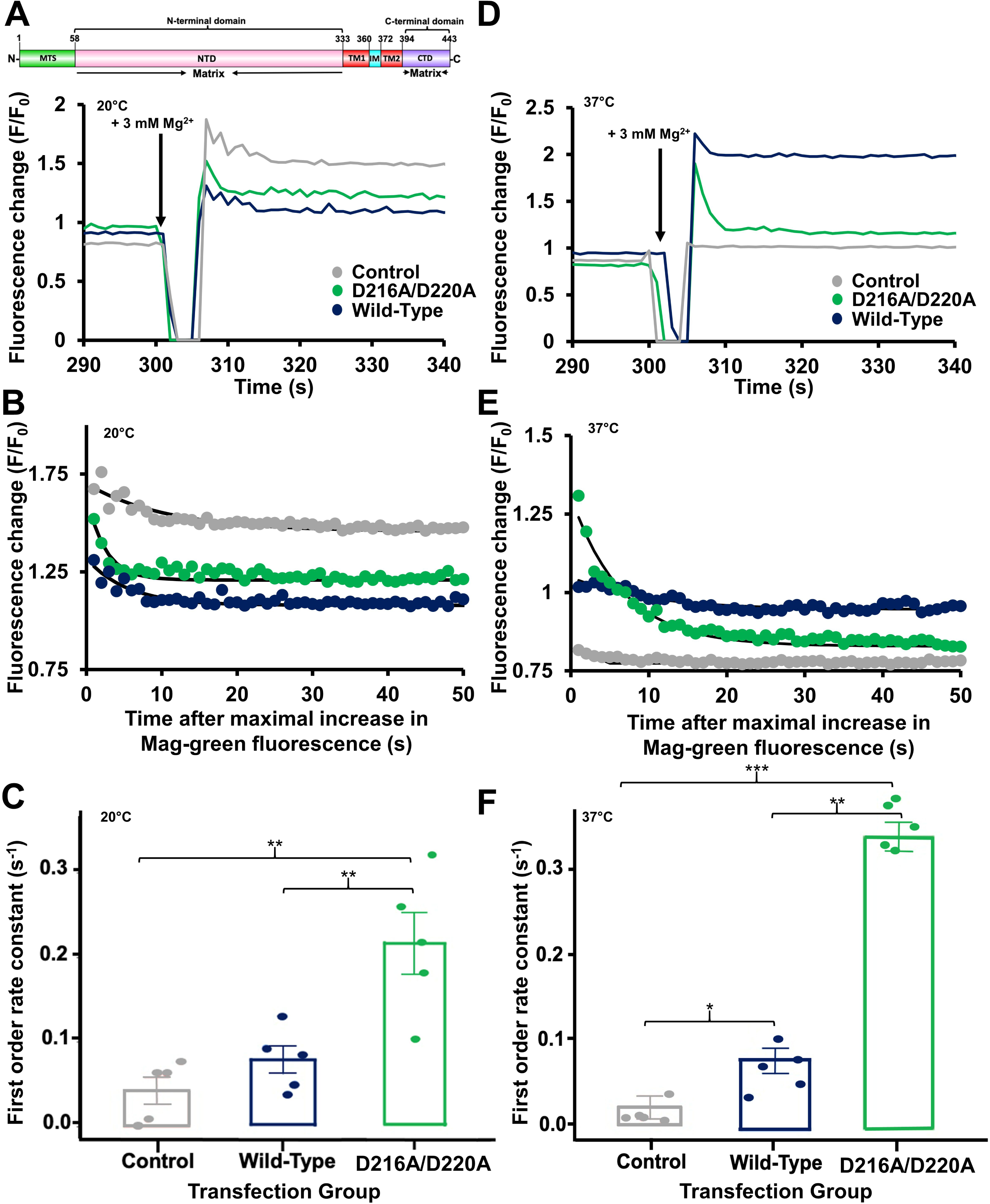
Mitochondrial Mg^2+^ uptake rates in HeLa cells overexpressing WT or D216A/D220A human MRS2. **(A)** Representative Mag-green fluorescence traces reporting relative change in intracellular/cytosolic Mg^2+^ before and after 3 mM MgCl_2_ addback to the extracellular bath at 300 s (black arrows) for control-, WT MRS2- and MRS2 D216A/D220A-transfected cells. **(B)** Representative single exponential fits to the Mag-green fluorescence decays after the 3 mM MgCl_2_ addback shown in (*A*), reporting intracellular Mg^2+^ clearance and taken as a measure of mitochondrial Mg^2+^ uptake. **(C)** One-way ANOVA followed by Tukey’s post-hoc comparison of mitochondrial Mg^2+^ uptake rates for control-, WT MRS2- and MRS2 D216A/D220A-transfected cells, where ** *p* < 0.01. **(D)** Representative Mag-green fluorescence traces reporting relative change in intracellular Mg^2+^ after 3 mM MgCl_2_ addback to the extracellular bath at 300 s (black arrows) for control-, WT MRS2- and MRS2 D216A/D220A-transfected cells. **(E)** Representative single exponential fits to the Mag-green fluorescence decays after the 3 mM MgCl_2_ addback shown in (*D*). **(F)** One-way ANOVA followed by Tukey’s post-hoc comparison of mitochondrial Mg^2+^ uptake rates for control-, WT MRS2- and MRS2 D216A/D220A-transfected cells, where * *p* < 0.05, \*\**p* < 0.01 and *** *p* < 0.001. In (*A-C*), data were acquired at 20 °C. In (*D-F*), data were acquired at 37°C. In (*D-F*), data were normalized as F/F_0_ where F is the Mag-green fluorescence at any time point and F_0_ is the mean 30 s baseline fluorescence prior to the addition of EDTA/digitonin, and control, WT and D216A/D220A data are coloured grey, black and green, respectively.

**Table S1:**
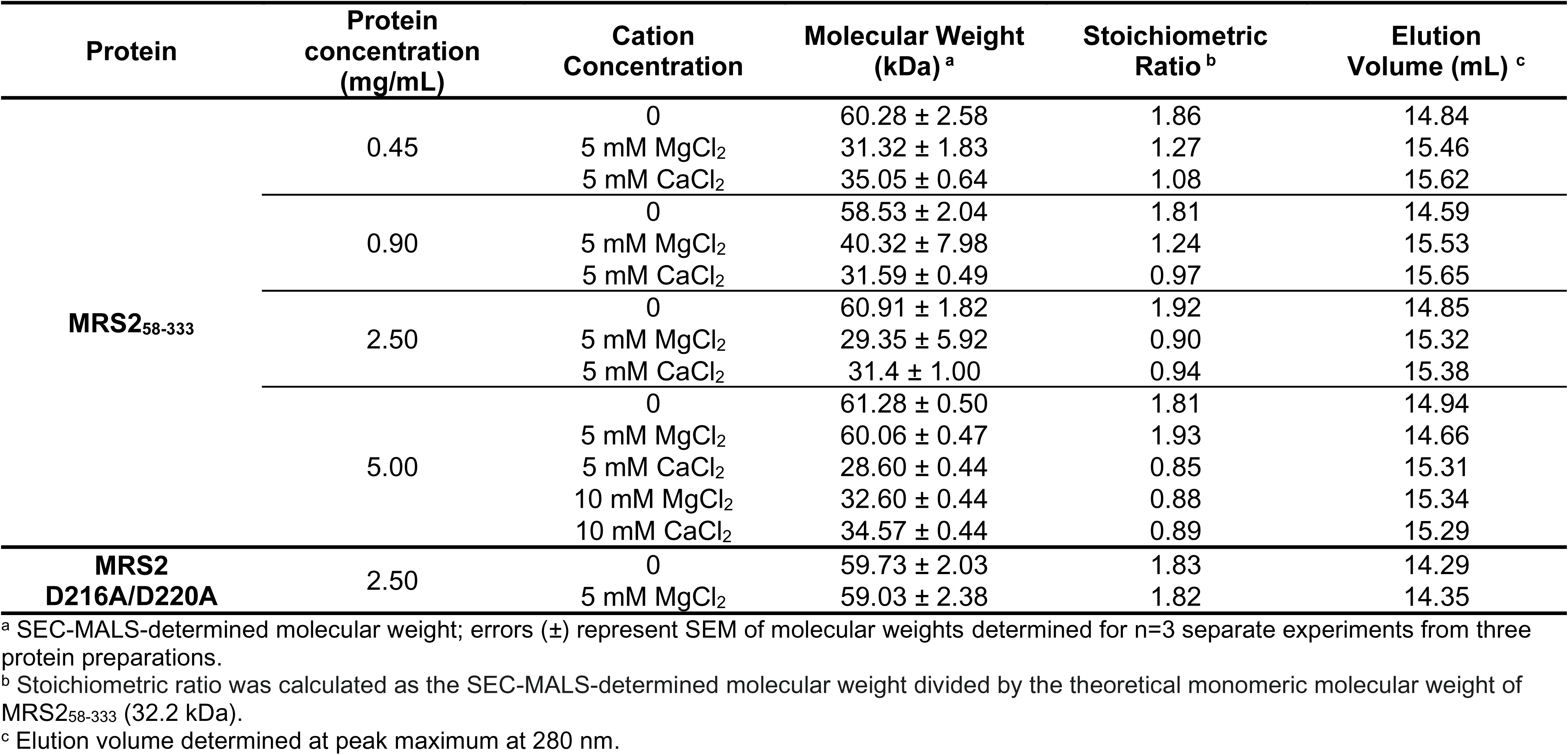
S**u**mmary **of SEC-MALS data for MRS_258-333_.**

**Table S2:** Summary of cumulants fits for MRS2_58-443_ and MRS2_58-333_.

| Protein | Temperature (°C) | Experimental Condition | Hydrodynamic Radius (nm)<sup>a</sup> | Experimental Condition<sup>b</sup> | Hydrodynamic Radius (nm)<sup>a</sup> |
| --- | --- | --- | --- | --- | --- |
| **MRS2<sub>58-443</sub>** | 20 | -MgCl<sub>2</sub> | 10.58 ± 1.04 | +MgCl<sub>2</sub> | 10.68 ± 0.84 |
|  |  | -CaCl<sub>2</sub> | 10.86 ± 1.25 | +CaCl<sub>2</sub> | 10.64 ± 0.95 |
|  |  | -CoCl<sub>2</sub> | 10.97 ± 1.38 | +CoCl<sub>2</sub> | 2.43 ± 1.33 |
|  | 37 | -MgCl<sub>2</sub> | 3.69 ± 0.50 | +MgCl<sub>2</sub> | 3.47 ± 0.90 |
|  |  | -CaCl<sub>2</sub> | 3.66 ± 0.78 | +CaCl<sub>2</sub> | 3.44 ± 1.25 |
|  |  | -CoCl<sub>2</sub> | 4.87 ± 0.95 | +CoCl<sub>2</sub> | 2.59 ± 1.04 |
| **MRS2<sub>58-333</sub>** | 20 | -MgCl<sub>2</sub> | 5.77 ± 0.88 | +MgCl<sub>2</sub> | 5.83 ± 1.14 |
|  |  | -CaCl<sub>2</sub> | 6.68 ± 0.85 | +CaCl<sub>2</sub> | 4.55 ± 0.96 |
|  |  | -CoCl<sub>2</sub> | 5.76 ± 0.50 | +CoCl<sub>2</sub> | 5.83 ± 0.49 |
|  | 37 | -MgCl<sub>2</sub> | 8.89 ± 0.46 | +MgCl<sub>2</sub> | 4.95 ± 0.64 |
|  |  | -CaCl<sub>2</sub> | 6.68 ± 0.70 | +CaCl<sub>2</sub> | 4.55 ± 0.96 |
|  |  | -CoCl<sub>2</sub> | 4.43 ± 0.53 | +CoCl<sub>2</sub> | 4.17 ± 0.77 |
| **MRS2 D216A/D220A** | 20 | -MgCl<sub>2</sub> | 5.69 ± 0.75 | +MgCl<sub>2</sub> | 5.67 ± 0.74 |

<sup>a</sup> Errors (±) are SEM from n=3 separate experiments from three protein preparations.
<sup>b</sup> +divalent cation indicates supplementation with 5 mM of the indicated metal salt.

**Table S3:**
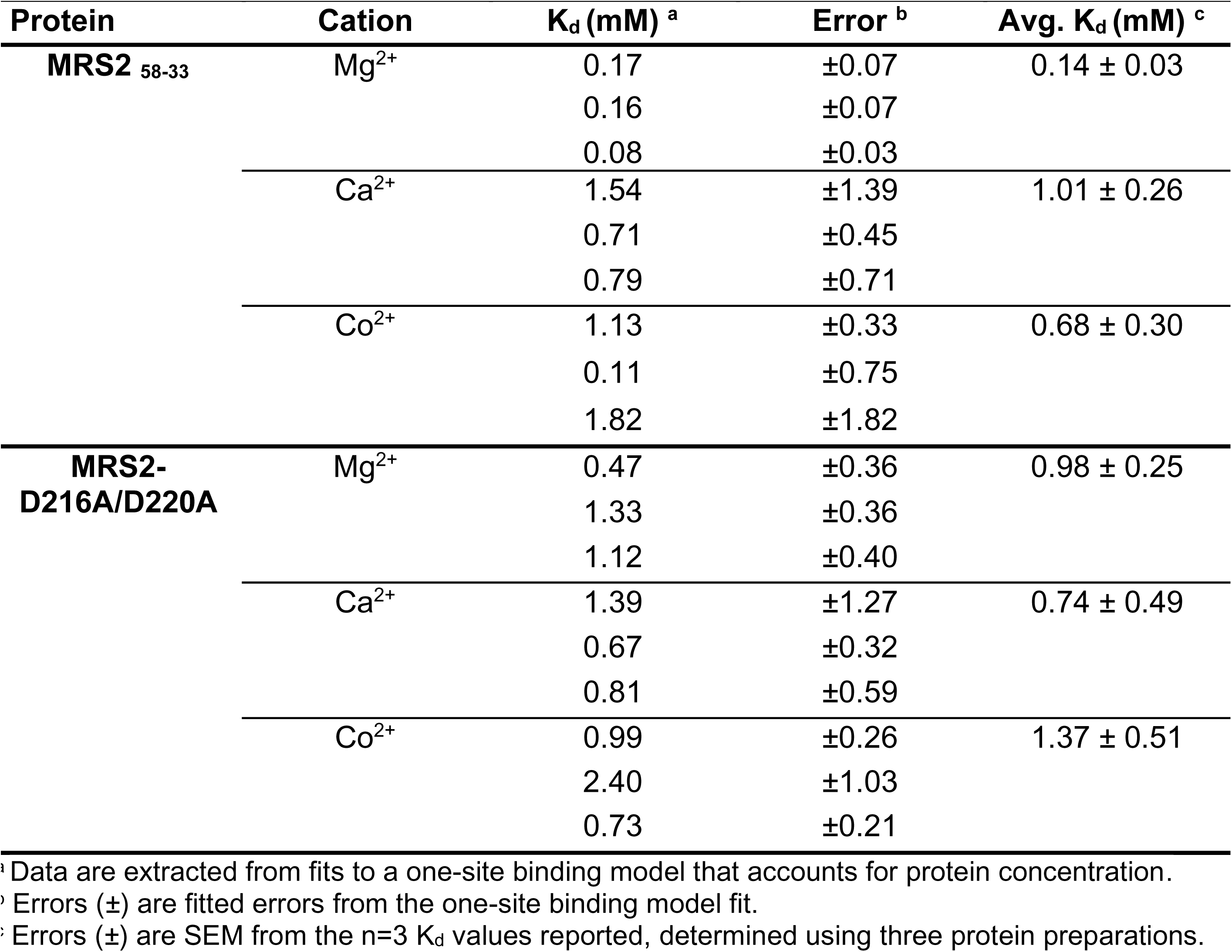
Summary of the fitted equilibrium dissociation constants (K_d_).

| Protein | Cation | $K_d$ (mM) <sup>a</sup> | Error <sup>b</sup> | Avg. $K_d$ (mM) <sup>c</sup> |
| --- | --- | --- | --- | --- |
| MRS2 <sub>58-33</sub> | Mg<sup>2+</sup> | 0.17 | ±0.07 | 0.14 ± 0.03 |
|  |  | 0.16 | ±0.07 |  |
|  |  | 0.08 | ±0.03 |  |
|  | Ca<sup>2+</sup> | 1.54 | ±1.39 | 1.01 ± 0.26 |
|  |  | 0.71 | ±0.45 |  |
|  |  | 0.79 | ±0.71 |  |
|  | Co<sup>2+</sup> | 1.13 | ±0.33 | 0.68 ± 0.30 |
|  |  | 0.11 | ±0.75 |  |
|  |  | 1.82 | ±1.82 |  |
| MRS2-<br>D216A/D220A | Mg<sup>2+</sup> | 0.47 | ±0.36 | 0.98 ± 0.25 |
|  |  | 1.33 | ±0.36 |  |
|  |  | 1.12 | ±0.40 |  |
|  | Ca<sup>2+</sup> | 1.39 | ±1.27 | 0.74 ± 0.49 |
|  |  | 0.67 | ±0.32 |  |
|  |  | 0.81 | ±0.59 |  |
|  | Co<sup>2+</sup> | 0.99 | ±0.26 | 1.37 ± 0.51 |
|  |  | 2.40 | ±1.03 |  |
|  |  | 0.73 | ±0.21 |  |

<sup>a</sup> Data are extracted from fits to a one-site binding model that accounts for protein concentration.
<sup>b</sup> Errors (±) are fitted errors from the one-site binding model fit.
<sup>c</sup> Errors (±) are SEM from the n=3 $K_d$ values reported, determined using three protein preparations.

